# Phylogenetic analyses for orthogroup-based classification of GDSL-type esterase/lipase (GELP) family in angiosperm representative species

**DOI:** 10.1101/2021.10.06.463335

**Authors:** Cenci Alberto, Concepción-Hernández Mairenys, Geert Angenon, Rouard Mathieu

**Author notes:** Corresponding author: Alberto Cenci.

## Abstract

GDSL-type esterase/lipase (GELP) enzymes have multiple functions in plants, spanning from developmental processes to the response to biotic and abiotic stresses. Genes encoding GELP belong to a large gene family with several tens to more than hundred members per angiosperm species.

Here, we applied iterative phylogenetic analyses to identify 10 main clusters subdivided into 44 expert-curated reference orthogroups (OGs) using three monocot and five dicot genomes. Our results show that some GELP OGs expanded while others were maintained as single copy genes.

This semi-automatic approach proves to be effective to characterize large gene families and provides a solid classification framework for the GELP members in angiosperms. The orthogroup-based reference will be useful to perform comparative studies, infer gene functions and better understand the evolutionary history of this gene family.

## Introduction

The first gene of the GDSL-type lipase/esterase (GELP) family was described by Upton and Buckley (Upton and Buckley 1995) in bacterial *Aeromonas* sp. with the amino acid motif of the active site (GDSL) distinguishable from one present in other Lipases (GxSxG). The GELP genes were later found widely distributed in all kingdoms. In plants, three main phylogenetic clusters were observed (Akoh et al. 2004), which have distinct exon/intron structures (Volokita et al. 2011). Through number comparison and phylogeny analysis of GELP family members, Volokita et al. (2011) showed a general trend of gene expansion in higher plants, which results in around 100 members in angiosperms.

GELP members were shown to be involved in a wide spectrum of functions, including hormone regulation, tissue development and biotic and abiotic stress responses (Volokita et al. 2011; Ding et al. 2019). The possible reactivity on different substrates was suggested for GELP enzymes (Huang et al. 2001). At present, not many GELP genes have been functionally characterized but it has been shown that they can play an important role in growth and development, plant immunity and biotic and abiotic stress responses (Ding et al. 2019; Shen et al. 2022).

It is anticipated that a number of crop specific transcriptomic studies will be conducted as was recently the case in Soybean (Su et al. 2020). An expert-curated orthogroup framework among GELP genes would be useful to transfer knowledge acquired on model species in non-model plant species, especially for such a large gene family. The elucidation of orthology relationships among genes should allow distinguishing orthogroups that underwent copy number expansion from those where only one or few copies per genome were maintained.

Functional annotation transfer requires an accurate identification of orthologous and paralogous sequences between genomes of different species and an expert-curated annotation of gene families appears to be a robust and reliable source of information. The identification of homologous relationships between genomes has been an active area of research, underlain by the orthology conjecture assuming that genes that diverged by speciation are functionally closer than those that diverged by duplication (Altenhoff et al. 2012). This hypothesis has been debated and recent analyses also reinforced the role of paralogs for functional prediction (Stamboulian et al. 2020). The relationship between two genes derived from a common ancestor by speciation is called orthology. When several species are considered, the relation of orthology can be described by orthogroups (OGs). Orthogroups are defined as the whole set of genes of a given sample of species descending from a single ancestral gene present in the most recent common ancestor of the considered species. Based on this definition, orthogroups determination is objective (based on phylogeny relationships) and relative (depending on the considered species). In other words, OG number and gene content will be different if the sampled species can be included in different classification ranks (e.g., angiosperms, dicots or rosids). As an evolutionary concept, homology is best inferred using phylogenetic analysis (Gabaldón 2008). However, reconstructing phylogenies for large gene families can be challenging due to a high number of homologous genes, presence of pseudogenes and possible annotation errors that introduce mistakes in sequence alignment and phylogeny inference. Automatic analyses were shown to produce inaccurate results on homologous clustering in the large family of NAC transcription factors (Cenci et al. 2014), although automatic methods continue to improve regularly (Glover et al. 2019; Emms and Kelly 2015).

Consequently, in order to obtain the most accurate phylogenetic relationships among genes of large families, a semi-automatic curation of genes and multi-step phylogenetic analyses were used in this study to establish orthogroups in large gene families such as transcription factors (NAC and GRAS) and Glycosyltransferase family 61 (Cenci et al. 2014; Cenci et al. 2018; Cenci and Rouard 2017). We further explored the gene exon-intron structure by orthogroups and functionally characterized GELP genes in the literature were also mapped to the catalog of orthogroups.

## Materials and Methods

### Sequence retrieval

All sequences annotated as “GDSL esterase/lipase” in *Amborella trichopoda* (GCA_000471905.1), *Phoenix dactylifera* (GCA_000413155.1), *Musa acuminata* v2 (GCA_904845865.1), *Oryza sativa* (GCA_001433935.1), *Coffea canephora* v1 (GCA_900059795.1), *Vitis vinifera* (GCA_000003745.2), *Prunus mume* v1 (GCA_000346735.1), *Theobroma cacao* v2 (GCF_000208745.1), and *Arabidopsis thaliana* (GCF_000001735.4), were retrieved from GenBank. BLASTp analysis (Altschul et al. 1990) was used to search misannotated genes using *V. vinifera* GDSL esterase/lipase genes as a query in the non-redundant protein sequence (nr) database of the nine sampled species. Sequences were visually inspected and, when necessary, corrections on structural annotation were performed (modified sequences are marked by “*”). Truncated sequences (missing part of the coding region due to genome sequence gaps) and probable pseudogenes having well-conserved protein sequences were tagged but maintained in the analyses.

### Phylogenetic analyses

Protein sequences were aligned with the MAFFT program (https://mafft.cbrc.jp) (Katoh and Standley, 2013) via the EMBL-EBI bioinformatics interface (Madeira et al. 2019) using default parameters. Sequence alignments were cleaned with GBlocks (Version 0.91b) (http://molevol.cmima.csic.es/castresana/Gblocks_server.html) (Castresana 2000). The cleaning was performed by allowing: (i) smaller final blocks, (ii) gap positions within the final blocks, and (iii) less strict flanking positions. Phylogenetic analyses were performed with PhyML (Guindon et al. 2009) available at http://www.phylogeny.fr (Dereeper et al. 2008) using an LG substitution model and an Approximate Likelihood-Ratio Test (aLRT). Unrooted phylogenetic trees were visualized with MEGA6 (Tamura et al. 2013).

### Orthogroups determination

OGs were established on phylogenetic trees including nine species and based on the last common ancestor of monocots and dicots (three and five species, respectively). *A. trichopoda* was not considered for OG definition and was considered as an outgroup. OGs were established according to the smallest clusters, including GELP sequences from both monocots and dicots.

### Sequence assignation to established OGs

Sequences OG assignment of GELP sequences from other species than the nine sampled in this study was based on BLASTp analysis. For each sequence, if the best five hits were consistently included in the same cluster/OG, the sequence was assigned to this cluster/OG.

## Results

All protein sequences annotated as GELP in nine angiosperm species representing major angiosperm branches (855 sequences) were manually inspected and, where necessary, gene structure corrections of the sequences were performed **(Supplementary material 1 and 2)**.

### Iterative phylogeny analysis of GELP sequences

In order to reduce the complexity of the analyses and to maintain angiosperm representation, 248 sequences from a subset of species were used to investigate the global GELP phylogeny: *A. trichopoda, P. dactylifera* and *V. vinifera* (66, 81 and 101, respectively) (**Table 1**). After the alignment filtering, only 102 positions remained. In the unrooted phylogenetic tree, a cluster (called Cluster 1) containing 14 sequences (respectively 2, 6 and 6 for the above cited species) was clearly separated from all the other ones (**Figure 1; Supplementary material 3, Tree 01; Supplementary material 4, Tree 01**). These sequences were removed from the dataset and phylogenetic analysis reiterated.

**Figure 1:**
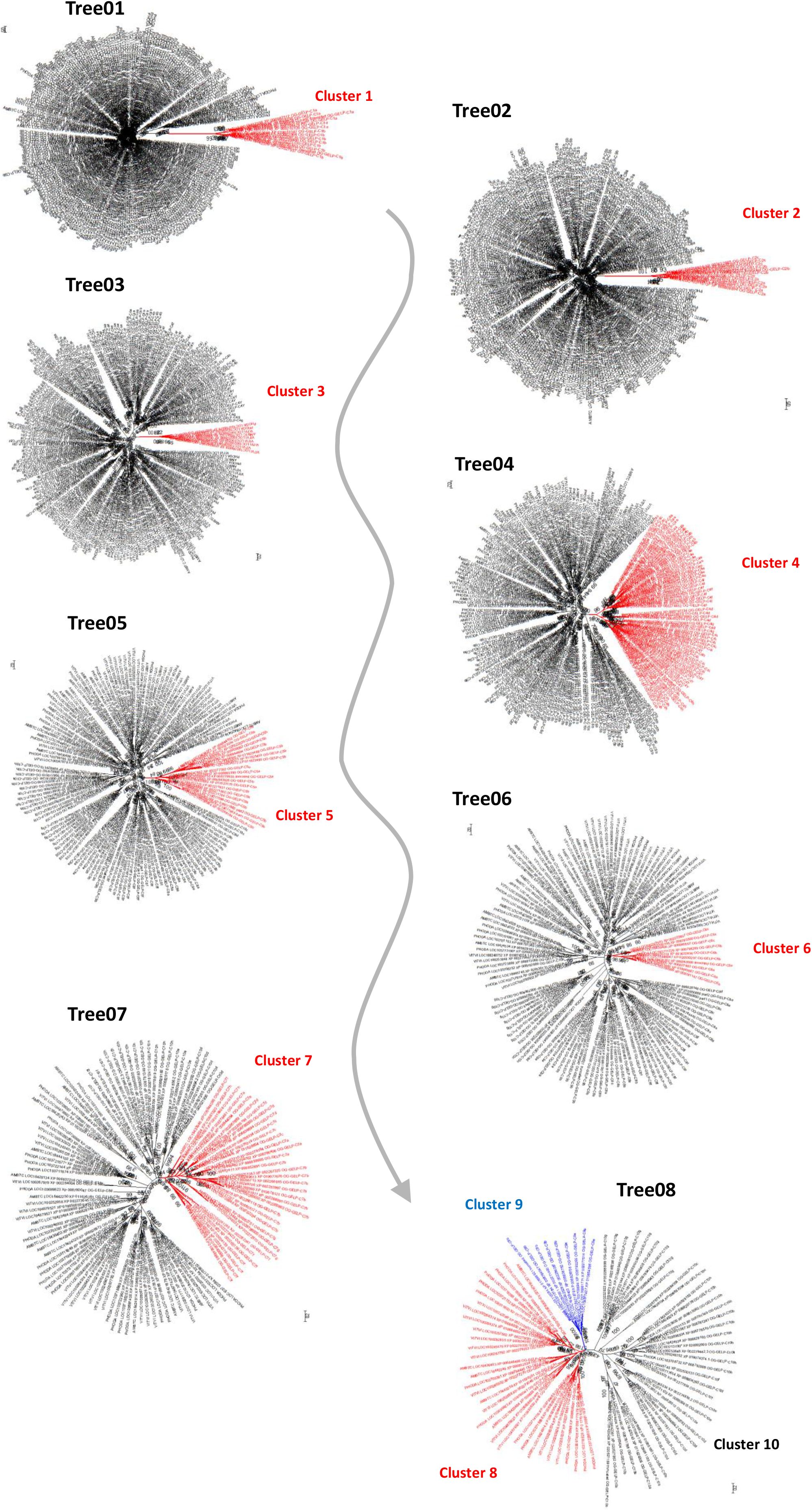
schematic illustration of the iterated strategy using unrooted phylogenetic trees.

**Table 1:** Iterated phylogenetic analyses. For each step, the number of analyzed sequences (*A. trichopoda-P. dactylifera-V. vinifera*), the number of aligned positions retained for phylogeny inference after alignment cleaning, the name of the tree in **Supplementary material 3** and **4,** the name assigned to the divergent cluster whose sequences were removed in following analyses (aLRT support of the cluster in the tree) and the number of sequences in the divergent cluster.

| Step | Sequence number | Positions | Tree name | Divergent cluster (aLRT support) | Sequences in cluster |
| --- | --- | --- | --- | --- | --- |
| 1 | 248 (66-81-101) | 102 | Tree01 | Cluster 1 (99) | 14 (2-6-6) |
| 2 | 234 (64-75-95) | 113 | Tree02 | Cluster 2 (100) | 12 (3-5-4) |
| 3 | 222 (61-70-91) | 135 | Tree03 | Cluster 3 (100) | 9 (1-2-6) |
| 4 | 213 (60-68-85) | 126 | Tree04 | Cluster 4 (96) | 65 (17-26-22) |
| 5 | 148 (43-42-63) | 179 | Tree05 | Cluster 5 (100) | 20 (10-2-8) |
| 6 | 128 (33-40-55) | 195 | Tree06 | Cluster 6 (98) | 10 (4-3-3) |
| 7 | 118 (29-37-52) | 204 | Tree07 | Cluster 7 (94) | 36 (8-6-22) |
| 8 | 82 (21-31-30) | 253 | Tree08 | Cluster 8 (90) | 33 (6-13-14) |
|  |  |  |  | Cluster 9 (100) | 9 (2-3-4) |
|  |  |  |  | Cluster 10 (97) | 40 (13-15-12) |

In the second round of phylogenetic analysis, 113 positions were retained after the masking step and used for gene tree reconstruction (**Table 1**). The phylogenetic tree was again characterized by a cluster (Cluster 2) well separated from the rest of the tree (**Figure 1; Supplementary material 3, Tree 02; Supplementary material 4, Tree 02**). Cluster 2 contains 12 sequences, respectively 3, 5 and 4 for the above cited species. The process was reiterated six more times (**Table 1**) until step 8 when the tree was split in three clusters (Cluster 8, 9, and 10) (**Figure 1; Supplementary material 3; Supplementary material 4**).

### Orthogroup definition

GELP sequences from *M. acuminata, O. sativa, P. mume, T. cacao, A. thaliana* and *C. canephora* **(Supplementary material 2)** were assigned to one of the ten clusters defined in the former analyses using BLASTp if the first five hits belong to the same cluster. All query sequences were assigned unequivocally to their respective cluster.

With the aim to establish monocot/dicot orthogroups (OGs) for the whole GELP family, the ten clusters, including the GELP sequences from *A. trichopoda*, the three monocots and five dicots, were submitted to a cluster-specific phylogenetic analysis. Orthogroups were established according to the smallest clusters, including sequences from both monocots and dicots.

#### Cluster 1

The phylogenetic tree obtained with the 47 sequences assigned to Cluster 1 (232 positions retained after masking) is composed of two main sub-clusters (**Supplementary material 3, TreeC1; Supplementary material 4, TreeC1**), each one containing at least one sequence for each of the nine species. In one sub-cluster, dicot and monocot sequences (5 and 10, respectively) are grouped separately and the *A. trichopoda* sequence has a basal position, consistent with the known phylogenetic relationships. The other sub-cluster is less resolved. If only branch support >0.9 are considered, one can observe some species-specific sub-clusters, isolated sequences (as the *A. trichopoda* one) and a dicot and a monocot sub-cluster. Based on these results, two orthogroups were established, OG-GELP-C1a and OG-GELP-C1b (**Table 2; Table 3**). We are aware that OG-GELP-C1b could be composed of sequences derived from two ancestor genes and consequently this OG could be split into two OGs.

**Table 2:** Overview of the orthogroups. Cluster name, Sequence number (*A. trichopoda, P. dactylifera, M. acuminata, O. sativa, V. vinifera, P. mume, T. cacao, A. thaliana, C. canephora*), the number of aligned positions retained for phylogeny after cleaning, the name of tree in Supplementary material 3 and Y.

| Cluster name | Sequence number | Positions | Tree name |
| --- | --- | --- | --- |
| Cluster 1 | 47 (2-6-5-7-6-4-5-9-3) | 231 | TreeC1 |
| Cluster 2 | 37 (3-5-4-4-4-4-4-4-5) | 217 | TreeC2 |
| Cluster 3 | 30 (1-2-3-2-6-8-1-5-2) | 254 | TreeC3 |
| Cluster 4 | 253 (17-26-35-54-22-23-25-25-27) | 169 | TreeC4 |
| Sub-cluster 4X | 50 (5-2-4-2-10-9-10-4-4) | 334 | TreeC4X |
| Sub-cluster 4Y | 149 (9-20-24-44-5-6-8-14-19) | 220 | TreeC4Y |
| Cluster 5 | 60 (10-2-0-0-8-12-10-11-7) | 284 | TreeC5 |
| Cluster 6 | 35 (4-3-4-3-3-5-6-4-3) | 319 | TreeC6 |
| Cluster 7 | 103 (8-6-8-7-22-14-15-15-8) | 240 | TreeC7 |
| Cluster 8 | 129 (6-13-12-18-14-13-11-33-9) | 262 | TreeC8 |
| Cluster 8X | 84 (3-6-7-10-10-7-6-29-6) | 275 | TreeC8X |
| Cluster 8X1 | 29 (2-1-2-3-3-2-4-8-4) | 295 | TreeC8X1 |
| Cluster 8X2 | 55 (1-5-5-7-7-5-2-21-2) | 279 | TreeC8X2 |
| Cluster 9 | 32 (2-3-4-3-4-3-3-5-5) | 312 | TreeC9 |
| Cluster 10 | 129 (13-15-20-24-12-15-11-9-10) | 213 | TreeC10 |
| Cluster 10X | 33 (1-4-5-5-4-3-3-4-4) | 334 | TreeC10X |
| Cluster 10Y | 37 (3-6-10-10-2-3-2-0-1) | 249 | TreeC10Y |

**Table 3:**
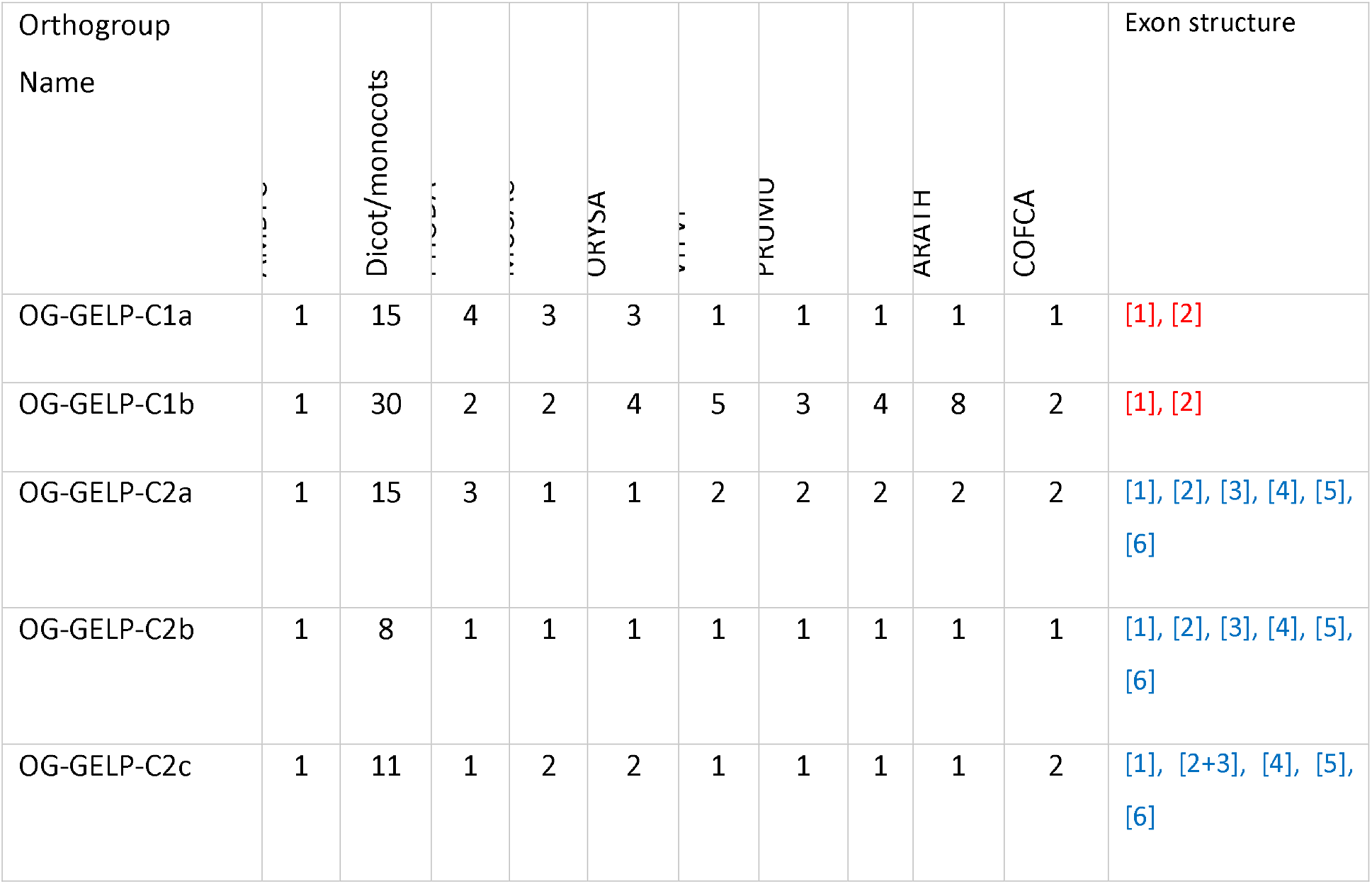
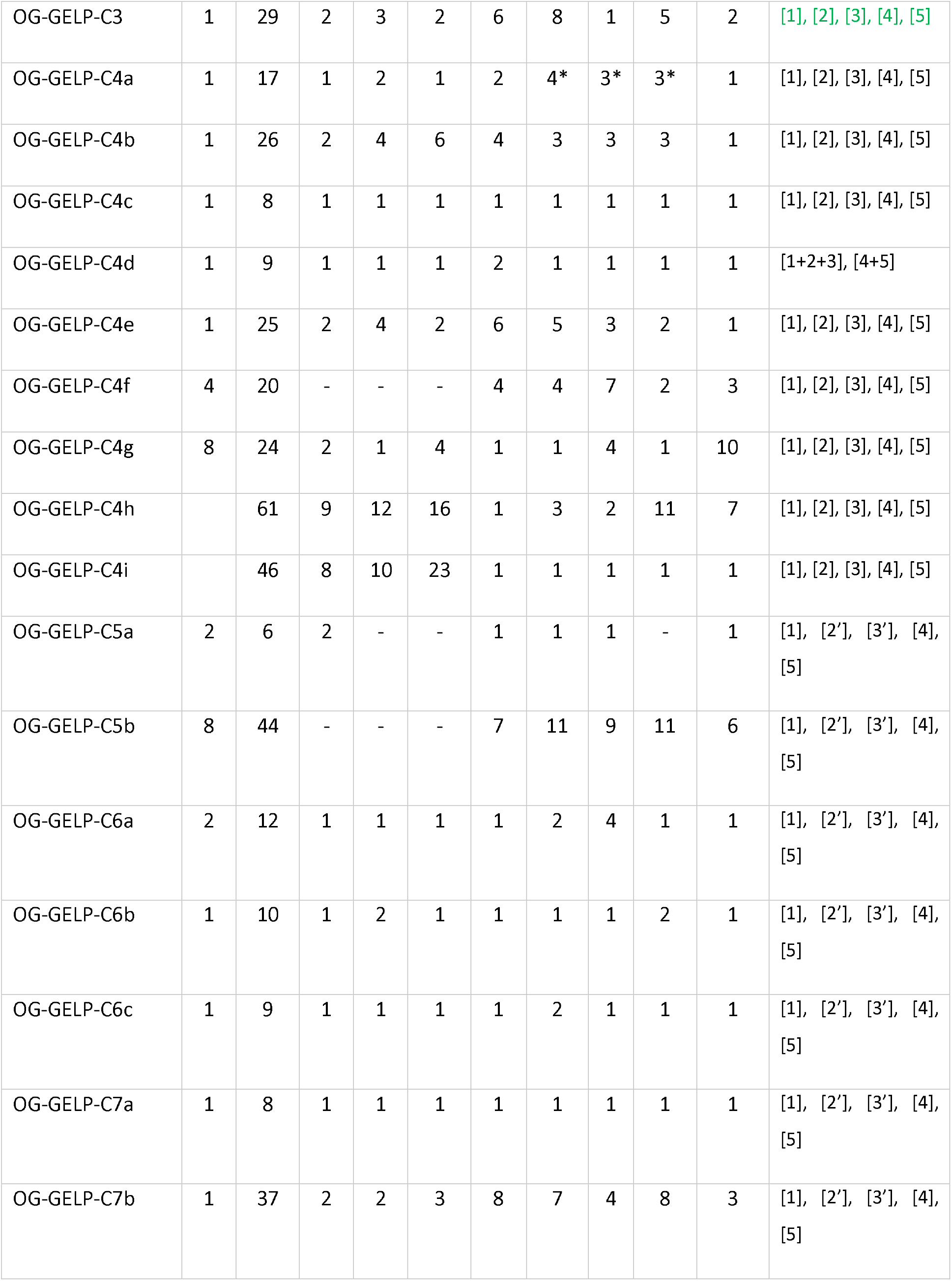
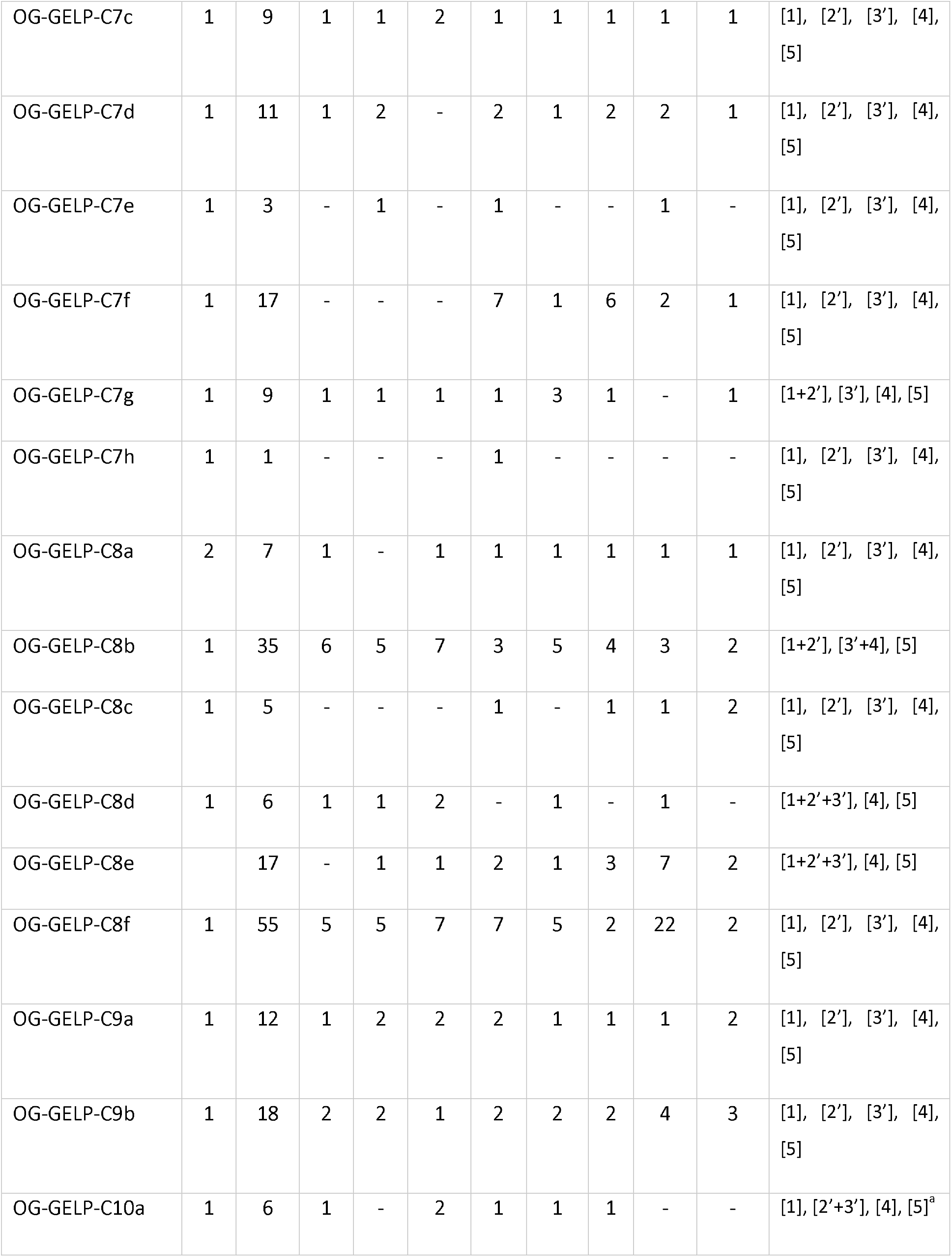
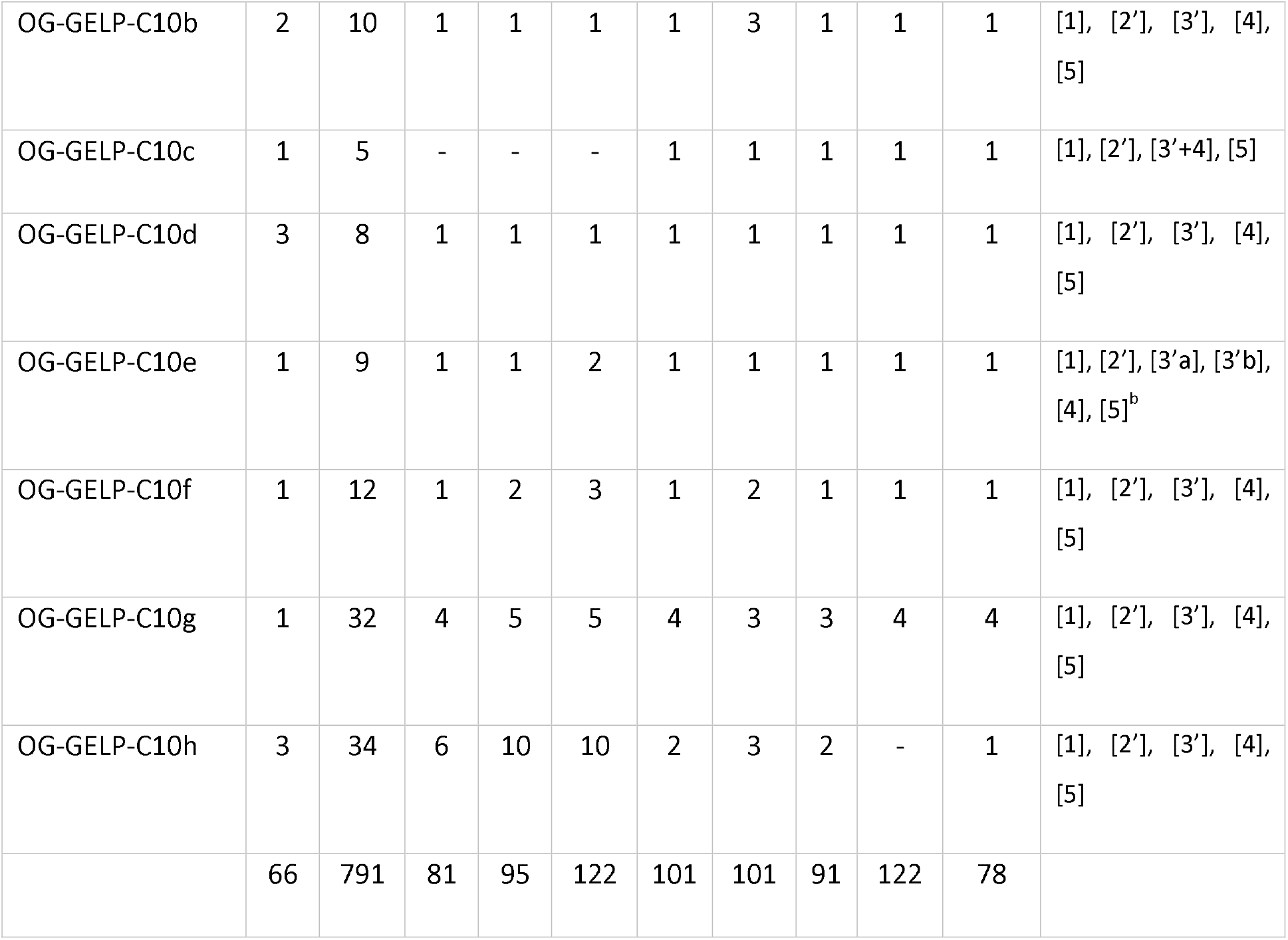
Per species sequence number assigned to the established GELP OGs. *Two or three sequences are alternative splicing versions where the first exon is chosen among 2 (or 3 in *P. mume*) independent regions. Independent exon structures are marked with different colors: red for genes in OGs derived from Cluster 1, blue for cluster 2, green for cluster 3 and black for the remaining clusters. Exons 2 and 3 having different junctions in sequences included in OGs derived from cluster 4 and clusters 5-10 are differentiated by a prime symbol (i.e., [2’] and [3’]).^a^ In dicots and *O. sativa* the structure is [1], [2’+3’+4], [5].^b^ in *A. trichopoda* exon [3’] is not split in two parts ([1], [2’], [3’], [4], [5])

#### Cluster 2

The phylogenetic tree obtained with 37 sequences assigned to Cluster 2 (207 positions) was structured in three main sub-clusters (**Supplementary material 3, Tree C02; Supplementary material 4, TreeC2**) with aLRT support spanning between 89 and 100, all containing at least one sequence of each one of the analyzed species. Also, in this case, some sub-clusters were less resolved. Three OGs were established with sequences included in Cluster 2: OG-GELP-C2a, -C2b and -C2c (**Table 2; Table 3**).

#### Cluster 3

The phylogenetic tree obtained with the 30 sequences assigned to the Cluster 3 (254 positions) was structured in two main clusters, one containing monocot species and the dicots and *A. trichopoda* (**Supplementary material 3, Tree C03; Supplementary material 4, TreeC3**). The sequences have been included in the OG-GELP-C3 (**Table 2; Table 3**).

#### Cluster 4

253 sequences from the nine sampled species were included in the Cluster 4. The phylogenetic analysis resulted in a tree containing two large unresolved sub-clusters and three strongly supported subclusters (aLRT>95), the last three all containing at least one sequence from each of the nine species and having a unique *A. trichopoda* sequence in basal positions (**Supplementary material 3, Tree C04; Supplementary material 4, TreeC4 and TreeC04b**). For each of the last three sub-clusters an orthogroup has been established: OG-GELP-C4a, -C4b and C4c (**Table 2; Table 3**). The two sub-clusters with poor resolution (hereafter called C4X and C4Y) were separately submitted to new phylogenetic analysis iterations.

##### Sub-cluster C4X

The phylogenetic tree built with the 50 GELP sequences included in Sub-cluster 4X was structured in two well supported sequence clusters (**Supplementary material 3, TreeC4X; Supplementary material 4, TreeC4X**). In one, both monocot and dicot sequences are represented, as well as a sequence from *A. trichopoda*. However, this cluster lacks a clear resolution, and the main nodes have poor support. The other cluster contains a sub-cluster of four *A. trichopoda* sequences with basal position and three highly supported clusters, containing at least one representative dicot sequence. No monocot sequences are present in this cluster. In several species, sequences included in both clusters are physically close: in *A. trichopoda* (all five sequences), in *P. mume* chromosome LG2 (LOC103321832/LOC103321833 and LOC103321831), in *V. vinifera* chromosome 1 (LOC100264465/LOC100259265/LOC100264440 and LOC109122650), in *T. cacao* chromosome 2 (LOC18609585/LOC18609586 and LOC18609588), in *C. canephora* chromosome 11 (CDP13237_Cc11_g15880 and CDP13238_Cc11_g15890). Based on these observations, we concluded that firstly, tandem duplication gave birth to the ancestors of two clusters before the split between *A. trichopoda* and the ancestor of monocot/dicot. Then, one copy was lost in monocot lineage, whereas the copy of dicot lineage underwent triplication before the asterid/rosid split. Two orthogroups were consequently established for the GELP sequences included in the Sub-cluster 4X: OG-GELP-4e and −4f (**Table 2; Table 3**).

##### Sub-cluster 4Y

The phylogenetic analysis of the Sub-cluster 4Y was performed on 220 positions retained in the alignment of 149 sequences (**Supplementary material 3, TreeC4Y; Supplementary material 4, TreeC4Y)**. A cluster containing 10 sequences from the nine species was sharply separated from the other sequences and the OG-GELP-C4d was established containing these sequences. In the remaining phylogenetic tree, all the *A. trichopoda* sequences are in the same cluster, whereas the other sequences formed three main clusters, all including both monocot and dicot species. Consequently, three OGs were established: OG-GELP-4g, -C4h and -C4i (**Table 2; Table 3**). In many cases, copy number expansion was observed both in species-specific and in taxonomically wider lineages.

#### Cluster 5

No sequences from *M. acuminata* or *O. sativa* are present in Cluster 5, making two *P. dactylifera* sequences the only representatives of monocots for our panel of species. The tree for Cluster 5 was built with 284 positions retained from an alignment of 60 sequences. Three main clusters are evident in the tree: the first *A. trichopoda* specific (8 sequences), the second with two tandemly located sequences from both *A. trichopoda* and *P. dactylifera* and one for all dicot species except *A. thaliana* and the third, the larger one, including 43 sequences from all dicots. The third cluster is structured in sub-clusters, some of them species-specific, composed of several tandem duplicated sequences. Even if the phylogeny was not completely resolved, two orthogroups were established based on second and third clusters, called OG-GELP-C5a and -C5b, respectively (**Table 2; Table 3**). The sub-cluster with eight *A. trichopoda* sequences, having six genes organized in tandem, was considered closer to the OG-GELP-C5b.

#### Cluster 6

The tree of cluster 6 was based on 319 aligned positions on 35 sequences. Three well-supported sub-clusters (aLRT>0.95) contain at least one representative sequence per species (**Supplementary material 3, TreeC6; Supplementary material 4, TreeC6**). Three orthogroups (OG-GELP-C6a, -C6b and -C6c) were established based on these sub-clusters (**Table 2; Table 3**).

#### Cluster 7

The tree of cluster 7 was based on 240 aligned positions on 103 sequences (**Supplementary material 3, TreeC7; Supplementary material 4, TreeC7**). Eight OGs were established based on main tree sub-clusters, each one containing at least a sequence of *A. trichopoda* and one from at least a monocot/dicot species but OG-GELP-7f missing monocots (**Table 2; Table 3**). OG-GELP-C7a contains a sequence for each species considered in this study. OG-GELP-C7b is the largest OG established in Cluster 7 and it is partially unresolved. Among the monocot/dicot evolution, repeated duplications took place after the separation from the *A. trichopoda* lineage, increasing the copy number in all the species studied here. In the OG-GELP-C7c, one sequence is included for each species, except for *O. sativa* which has two sequences, one with a basal position in the sub-cluster containing the other sequences included in the OG. OG-GELP-C7d is missing sequences from *O. sativa* and two *M. acuminata* sequences with isolated basal position were tentatively included in the OG with poor support. OG-GELP-C7e contains only a few sequences belonging to both monocots and dicots as well as to *A. trichopoda*. In *V. vinifera* and *T. cacao*, the gene underwent gene copy amplification. Only sequences from the dicot species were included in the OG-GELP-C7f. An *A. trichopoda* sequence (AMBTC_LOC18433946) appears close to this OG, however the aLRT support was very low and its association with the OG-GELP-C7f should be considered uncertain. OG-GELP-C7g contains sequences from all the species except *A. thaliana*. BLASTp analyses indicated that all species in the Brassicales order are missing members of the OG-GELP-C7g. One sequence from *A. trichopoda* and one from *V. vinifera* were associated in a two-sequence highly supported cluster. BLASTp analyses in the non-redundant protein database found only one sequence of *Nelumbo nucifera* (XP_010258913) close to the one in this cluster. According to the phylogenetic evidence, the OG-GELP-C7h was established including only these two sequences.

#### Cluster 8

Cluster 8 tree was based on 261 positions on 129 sequences (**Supplementary material 3, TreeC8; Supplementary material 4, TreeC8**). OG-GELP-C8a (7 sequences) and -C8b (35 sequences) were established according to two strongly supported clusters (aLRT=0.99) (**Supplementary material 3, TreeC8b; Supplementary material 4, TreeC8b**), both containing sequences from monocots and dicots as well as one or two sequences from *A. trichopoda* (**Table 2; Table 3**). The 84 sequences contained in the remaining poorly resolved cluster C8X were used to build a tree based on 275 positions. This tree was structured on two main clusters (aLRT=0.99), called C8X1 (29 sequences) and C8X2 (55 sequences) (**Supplementary material 3, TreeC8X; Supplementary material 4, TreeC8X**). For each sub-cluster the phylogenetic analysis was iterated. The tree obtained with the C8X1 sequences was based on 295 positions (**Supplementary material 3, TreeC8X1; Supplementary material 4, TreeC8X1**). Three OGs were defined with these sequences (**Table 2; Table 3**): OG-GELP-C8c, with high branch support was missing monocot sequences and contains an *A. trichopoda* sequence; two other OGs were established on clusters supported at 0.90 (OG-GELP-C8d) and 0.96 (OG-GELP-C8e). These OGs contain both monocot and dicot sequences. An *A. trichopoda* sequence has an intermediate position (basal to the OG-GDSM-C8d cluster but with support lower than 0.9). The tree obtained with the 55 C8X2 sequences (279 positions) was structured in two large clusters, one specific to monocots and the other specific to dicots (**Figure 2; Supplementary material 3, TreeC8X2**). Consequently, all these sequences were included in a unique OG, named OG-GELP-C8f (**Table 2; Table 3**). In this OG, copy amplification occurred, increasing the gene copy number. In dicots, the amplification appears more recent than the radiation of the analyzed species (21 *A. thaliana* sequences are present, organized in three main species-specific clusters, whereas seven sequences are in a *V. vinifera* specific cluster). On the contrary, in monocots, gene amplification took place before the radiation of Commelinidae, generating at least four ancestral genes still present in the analyzed species (**Figure 2**).

**Figure 2:**
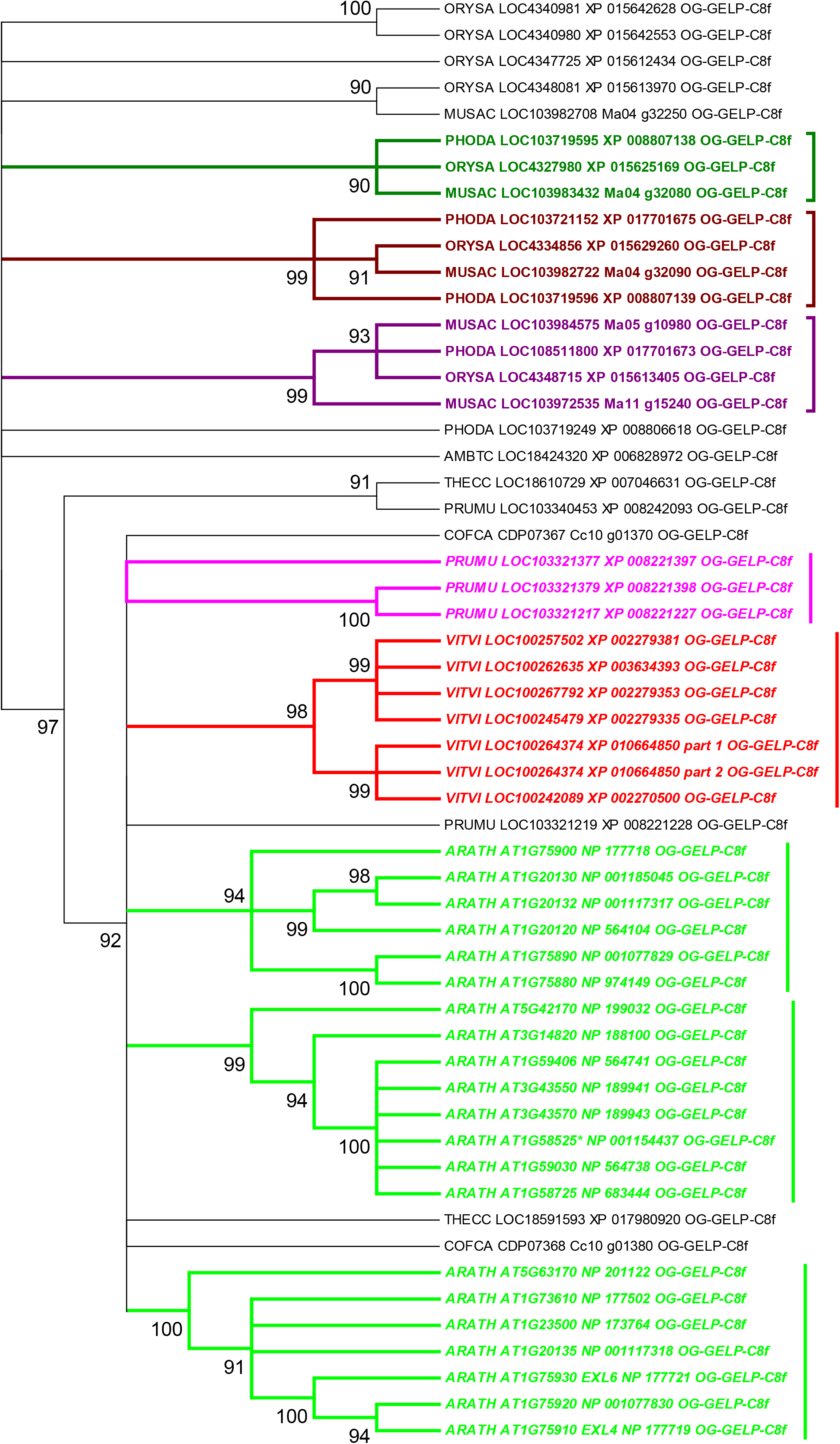
Unrooted phylogenetic tree built with sequences assigned to the OG-GELP-C8f. Nodes with aLRT support lower than 0.9 are collapsed. Colored sub-clusters include sequences originated by duplications that occurred before species radiation (in monocot subtree, square brackets) or species-specific duplication (in dicot subtree, vertical line).

#### Cluster 9

The Cluster 9 tree was based on 312 positions on 32 sequences. Sequences were distributed in two main clusters, both containing sequences from monocots, dicots and *A. trichopoda* (**Supplementary material 3, TreeC9; Supplementary material 4, TreeC9**). Two OGs were established: OG-GELP-C9a and OG-GELP-C9b (**Table 2; Table 3**). In the second cluster a dicot specific duplication can be inferred from the tree.

#### Cluster 10

The 129 sequences included in Cluster 10 were used to build a phylogenetic tree based on 210 aligned positions. Eight well-supported clusters (aLRT > 0.95) can be distinguished **(Supplementary material 3, TreeC10; Supplementary material 4, TreeC10)**. Six of them have a low number of sequences per species (3 or less) and contain at least one *A. trichopoda* sequence and representative sequences of both monocots and dicots (except one that is missing monocots). Based on these clusters, six OGs were established (from OG-GELP-C10a to OG-GELP-C10f in **Table 3**). Two other large sub-clusters (aLRT>0.97) contain a higher number of sequences per species and are poorly resolved (Cluster C10X and C10Y). Both sub-clusters are not organized in two clearly separated monocot and dicot specific sub-clusters. When the 32 sequences of the C10X sub-cluster were used to build a new phylogenetic tree (based on 334 aligned positions), two main clusters were observed (aLRT=0.91), one containing all the 14 monocot sequences, the other including one *A. trichopoda* and 17 dicot sequences **(Supplementary material 3, TreeC10X; Supplementary material 4, TreeC10X)**. In the OG-GELP-C10g obtained from the C10X sequences (**Table 3**), gene copy amplification before dicot species radiation was observed, whereas in monocots gene copy amplification was mainly lineage specific **(Figure 2)**. Finally, even if the phylogenetic relationships among the 39 sequences included in the C10Y sub-cluster could not be completely resolved, two main clusters were observed, including the one with all monocots and the other with most of the dicots **(Supplementary material 3, TreeC10Y; Supplementary material 4, TreeC10Y)**. The OG-GELP-C10h was tentatively established based on this cluster (**Table 3**).

### Gene structure of GELP genes

For each OG (44 in total for 855 genes) the exon structure is conserved, with few exceptions observed. When the exon structures were compared between OGs, five main structures were observed (Table 3). 1) Sequences in OGs derived from Cluster 1 have only 2 exons with aligned exon junctions, the second exon including more than 85% of the gene length. 2) The sequences assigned to three OGs derived from Cluster 2 (OG-GELP-C2a, -C2b and -C2c) have similar structures with aligned exon junctions. Sequences in OG-GELP-C2a and -C2b have 6 exons, whereas the ones in OG-GELP-C2c have 5, due to the fusion of second and third exons. 3) The sequences in OG-GELP-C3 have 5 exons with junctions located in different positions when compared to the genes included in the other OGs. 4) Sequences in 8 of the 9 OGs derived from Cluster 4 have 5 exons with aligned junctions, the remaining one, OG-GELP-C4d, being composed of only 2 exons, obtained by the fusion of 3 and 2 exons, respectively. 5) Sequences in all other OGs share a basic 5-exon structure, like the one observed for the OGs derived from Cluster 4, however the junction between exon 2 and 3 is different. According to specific OGs, the number of exons can be reduced by fusion of two or more exons or, in the case of OG-GELP-C10e, increased by the insertion of an intron in the exon 3. Some dicot sequences of OG-GELP-C4a have two or three alternative first exons (Table 3).

### Functionally characterized GELP genes

Few GELP genes have been functionally characterized (**Table 4**). The list of Ding et al. (Ding et al. 2019) was integrated with some other GELP genes. Classification of these genes (sequences reported in **Supplementary material 5**) was performed by BLASTp on a database containing the GELP genes of this study (**Supplementary material 2**). For each sequence not belonging to the nine studied species, the assignment was claimed if at least the first five hits consistently belong to the same OG.

**Table 4.**
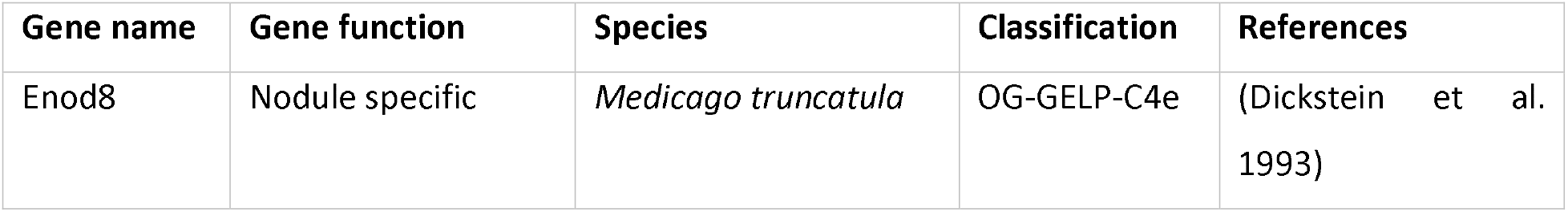
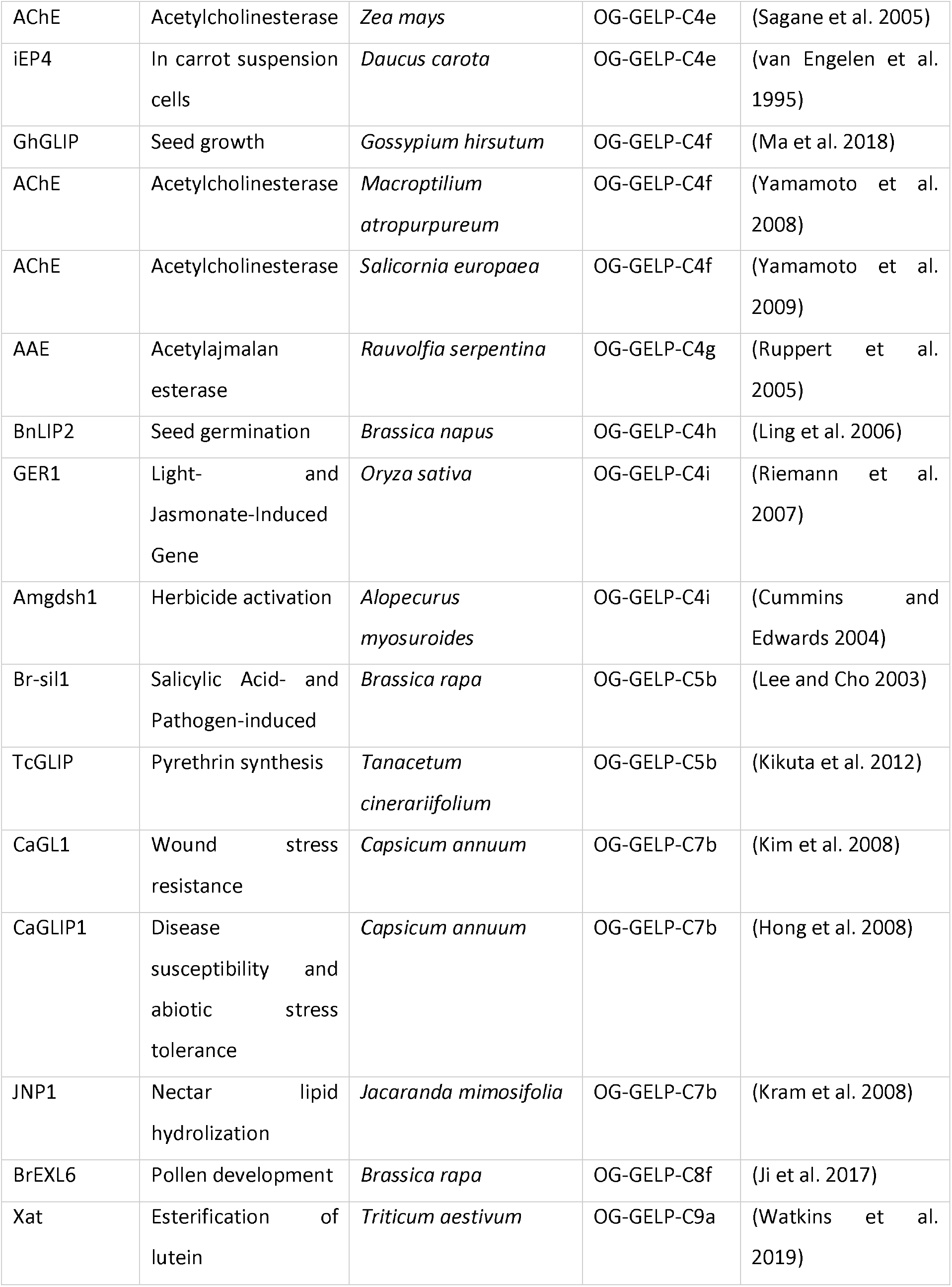
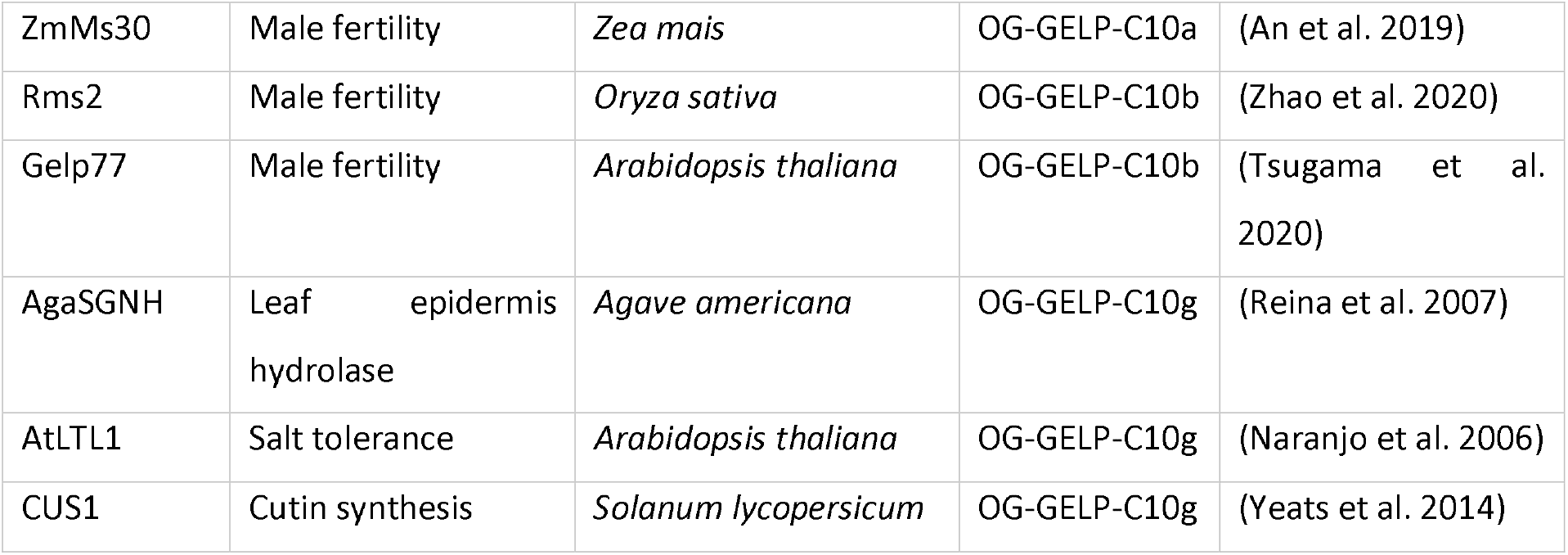
Classification of described GELP genes.

## Discussion

### An iterative phylogenetic approach for large gene families

The reconstruction of phylogenetic relationships among genes belonging to families with many members is challenged by the number of sequences. Calculation time increases exponentially with the number of sequences, and at the same time, the sequence alignment process could be challenged by the sequence divergence. Moreover, gene sequence inaccuracies produced during automatic annotations could introduce biases in the alignments (Gottin et al., 2021). Sequence accuracy can be improved by inspecting sequences, especially when parts of them appear poorly aligned to the closest homologs. In our study, genomic sequences were manually inspected and when needed, alternative annotations manually determined based on similarity with homologs. Pseudogene sequences were identified if evidence of mutations was found that were bypassed by automatic annotation by exon/intron structure adjustment.

To cope with the difficulty due to the large number of sequences, we adopted strategies aimed to reduce the number of analyzed sequences. Even if genes from nine species were considered in this study (with more than 850 GELP sequences) to represent major angiosperm branches, only the sequences from three species were analyzed in the global phylogenetic reconstruction: one representing monocots (*P. dactylifera*), one representing dicots (*V. vinifera*) and *A. trichopoda*, which has phylogenetic position basal to the monocot/dicot lineage split. The representative species for monocot and dicot lineages were chosen based on the lower number of whole genome duplication in their genome history.

The second strategy we used to reduce the difficulties induced by the sequence number on alignment and phylogenetic accuracy was an iterative approach. In fact, the larger the number of analyzed sequences, the lower the number of positions for phylogenetic inference remained after removing poorly aligned columns. A clear negative correlation between the number of analyzed sequences and aligned positions can be inferred from **Table 1** and **Table 2.** Phylogenetic trees were cleaved in main clusters with very strong support and the phylogenetic process (alignment, masking and phylogenetic inference) was repeated for each reduced dataset. This zoom-in approach allows recovering informative positions discarded in more complex alignments and provides better phylogenetic resolution.

Eight analysis iterations on the three sampled species (248 sequences of the whole genomic GELP set) resulted in 10 clusters (**Figure 1**). Except for few gene specific peculiarities, exon/intron structures were consistent with this first level classification (**Table 3**). Clusters 1, 2, 3, have completely independent intron positions. Exon structures of sequences in Clusters 4 and 5-10 are different from the ones in the former clusters and are partially consistent, the only difference being the insertion point of the second intron.

### Comparison with previous analyses

The iterative phylogenetic reconstruction of the GELP family obtained in this study is consistent with former phylogeny reconstructions based on the complete gene set of GELP genes on *O. sativa, A. thaliana* and *P. mume* (Chepyshko et al. 2012; Volokita et al. 2011; Lai et al. 2017; Cao et al. 2018). However, the sequences from our Cluster 1 were not considered in these studies. Indeed, these sequences have been annotated as GDSL esterase/lipases in several species but, even if sharing some degree of homology with other sequences, they encode for proteins not bearing the GDSL motif and are annotated as Alpha/beta-Hydrolase in *A. thaliana*. Consequently, the separated clustering of these sequences is not surprising. The genes included in Cluster 1 are composed of only two exons.

Sequences in Cluster 2 were not considered by Volokita et al. (2011) nor by Lai et al. (2017) and even if counted among rice GELP genes by Chepyshko et al. (2012), the four *O. sativa* sequences in the Cluster 2 were excluded from their phylogenetic analysis. In the study of Cao et al. (2018) focused on Rosaceae GELP family, *P. mume* genes included in Cluster 2 are part of Subfamily A. Cluster 2 sequences have GSSI or GDSI sequences in the place of the typical GDSL motif and the genes have a typical structure in 6 exons.

Sequences in Cluster 3 are characterized by the GDSY sequence at the place of the GDSL motif. This cluster includes the sequences in Clade C in Volokita et al. (2011), Clade IVa in Lai et al. (2017), Clade II in Chepyshko et al. (2012), and Subfamily C in Cao et al. (2018). The genes included in the Cluster 3 have 5 exons.

The fourth phylogenetic iteration clearly separated sequences in Cluster 4 that are included in Clade B in Volokita et al. (2011), Clades IVb and IVc in Lai et al. (2017), Clade I in Chepyshko et al. (2012), and Subfamily D in Cao et al. (2018). Like the GELP genes on Cluster 3, genes included in Cluster 4 have 5 exons, but their splicing positions are different.

Finally, the Clades A1 and A2 on Volokita et al. (2011) as well as Clades I + II and Clade III, in Lai et al. (2017) correspond, in our reconstruction, to Clusters 6-10 and 5, respectively. *O. sativa* sequences in Clade III and IV on Chepyshko et al. (2012) correspond to Cluster 5-10. The *P. mume* sequences included in subfamilies B and E-J of the Rosaceae GELP (Cao et al. 2018) are represented in Clusters 5-10. The five-exon structure observed for the genes in Clusters 5-10 is similar to that of Cluster 4 genes, with the exception of the second splicing site which has a different position.

Few inconsistencies with previously inferred phylogenies were observed in lower rank (more recent clades) of our phylogeny and we consider that the larger species sample size along with the lower sequence divergence in our iterated phylogenetic analyses provides a more accurate phylogenetic inference and granular classification of the GELP gene family. Based on the monocot/dicot split, 44 OGs were established, most of them containing representatives of all species (**Table 3**).

### An expert curated dataset to facilitate classification

The dataset on which our analyses were conducted (**Supplementary material 2**) includes the 786 monocot/dicot GELP sequences assigned to the 44 OGs as well as the 66 *A. trichopoda* sequences useful as outgroups in OG establishment. In the multifasta file, each sequence name begins with the five digit species code and ends with the name of the respective OG. This file could be used as a database for BLASTp analysis to rapidly assign to one of the 44 OGs here established the whole genome sets of GELP genes as well as specific GELP genes from any monocot/dicot species.

As an example, the GELP genes listed by Ding et al. (2019) in Table 1 were assigned to their respective OG (**Table 4; Supplementary material 5**). Along with consistency between OG classification and known function (e.g., ZmMs30, Rms2 and Gelp77, respectively; included in OG-GELP-10a and −10b and involved in male fertility in *Z. mays, O. sativa* and *A. thaliana*, respectively), the classification of these and other GELP characterized genes allows highlighting phylogenetic proximity between genes involved in apparently different functions or the phylogenetic distance between genes involved in similar functions. The *A. thaliana* gene FXG1 (α-Fucosidase) (de la Torre et al. 2002) belongs to the same OG-GELP-C4e as the *Medicago sativa* gene coding for ENOD8 protein (present in root nodules) (Pringle and Dickstein 2004) and the *Daucus carota* gene coding for the EP4 glycoprotein observed in the cell wall in seedling roots (van Engelen et al. 1995) as well as to the genes coding for *Zea mays* and *A.thaliana* acetylcholine esterases (Sagane et al. 2005; Muralidharan et al. 2013). Conversely, the acetylcholine esterase genes isolated in *Macroptilium atropurpureum* and in *Salicornia europaea* (Yamamoto et al. 2008;Yamamoto et al. 2009) belong to the phylogenetically close but different OG-GELP-C4f, which also includes the *Gossypium hirsutum* gene GhGLIP shown to be involved in seed development (Ma et al. 2018). According to the OG classification, the five GELP genes (SFAR1-5) shown to be involved in *A. thaliana* fatty acid degradation during the seed germination (Chen et al. 2012) belong to different OGs (OG-GELP-C4a, -C8e, -C8b, -C4d, and -C7b) that do not have close phylogenetic relationships.

### Gene duplicates

Copy number amplification appears as a global trend in the evolution of the GELP family, however our results indicated that if in some OGs the copy number increased after the monocot/dicot split, in some others, even phylogenetically close OGs, the copy number did not change and was limited to one sequence per species (**Table 3** and **Supplementary material 4**). The copy number expansion was due to gene duplications, both local (tandem duplication) or distal (possibly generated by whole genome duplications). The observed duplications are either lineage specific or having taken place before species radiation (e.g., OG-GELP-C8f and -C4f, **Figure 2**). The extreme cases of GELP gene amplification are the 22 *A. thaliana* genes in the OG-GELP-C8f. These differences suggest that some GELP ancestor genes evolved under different evolutionary forces, possibly due to their implication in different functions. For each OG, the detailed duplication history is represented in respective phylogenetic trees (**Supplementary material 3; Supplementary material 4**). Tandem duplications contributed to the gene copy amplification, after the monocot/dicot split, but also before this evolutionary milestone, as testified by the tandem position of representatives of OG-GELP-C8a and -C8c, in *A. trichopoda, C. canephora, A. thaliana* and *T. cacao* or of OG-GELP-C4e and -C4f in *A. trichopoda, P. mume, V. vinifera, C. canephora,* and *T. cacao*.

### Functional transfer in large gene families

All the genes included in an OG derived from the same ancestor gene that was present in the common ancestor species of monocot and dicot. The OG framework should facilitate the functional annotation transfer between species, highlighting the phylogenetically closest genes. However, the functional transfer of genes belonging to OGs with a high number of members should be carefully evaluated. In fact, the persistence of multiple copies derived from a single ancestor gene could be the consequence of sub- or neo-functionalization that occurred among phylogenetically close genes and independently evolved new functions are likely different. The OG framework inferred in the present study provides detailed information allowing the discrimination between gene lineages that underwent diversification from the ones that appears stable from an evolutive point of view. The nine-species GELP sequence set (**Supplementary material 2**) could also be used as a verified kernel for an orthogroup inference of the GELP family on a large number of species.

## Acknowledgements

This work was supported by the Belgian Directorate-General for Development Cooperation (DGDC) and the CGIAR Research Project Roots, Tubers and Bananas (CRP-RTB) and by the VLIR-UOS Research Initiatives Program ZEIN2016RIP33. The authors are grateful to Rachel Chase for careful English editing.

## Supplementary Material

### Supplementary material 1.xlsx

GELP sequence names analyzed in this study. Sequences are grouped based on Orthogroup classification. For *O. sativa* and *P. mume* the name assigned in previously published studies is reported (Chepyshko *et al.* 2012; Cao *et al.* 2018).

### Supplementary Material 2.fa

Sequences of 855 GELP genes used in this study. In the sequence names, the first five digits indicate from which of the nine species the sequence was retrieved, the last three digits identify the OG to which the sequence was assigned.

### Supplementary Material 3.txt

Phylogenetic trees (Newick format) inferred in this study. Tree names refer to **Table 1** and **Table 2.**

### Supplementary Material 4.pptx

Graphic representations of phylogenetic trees inferred in this study. In tree radial representation different colors highlight well supported clusters on which the tree was divided. In rectangular tree representations different branch colors indicate the Orthogroup (OG) defined based on phylogenetic tree, whereas sequence colors (where applicable) indicate sub-clusters grouping monocot/dicot sequences, highlighting duplications occurred after the monocot/dicot split or species-specific gene clusters. In rectangular tree representations the right dendrogram is based on nodes supported with aLRT>90.

### Supplementary Material 5.fa

Sequences of 19 GELP genes functionally characterized and belonging to species that were not sampled in our study.

