## Supplementary file 4 for "Phylogenetic analyses for orthogroup-based classification of GDSL-type esterase/lipase (GELP) family in angiosperm representative species"

### Slide 1
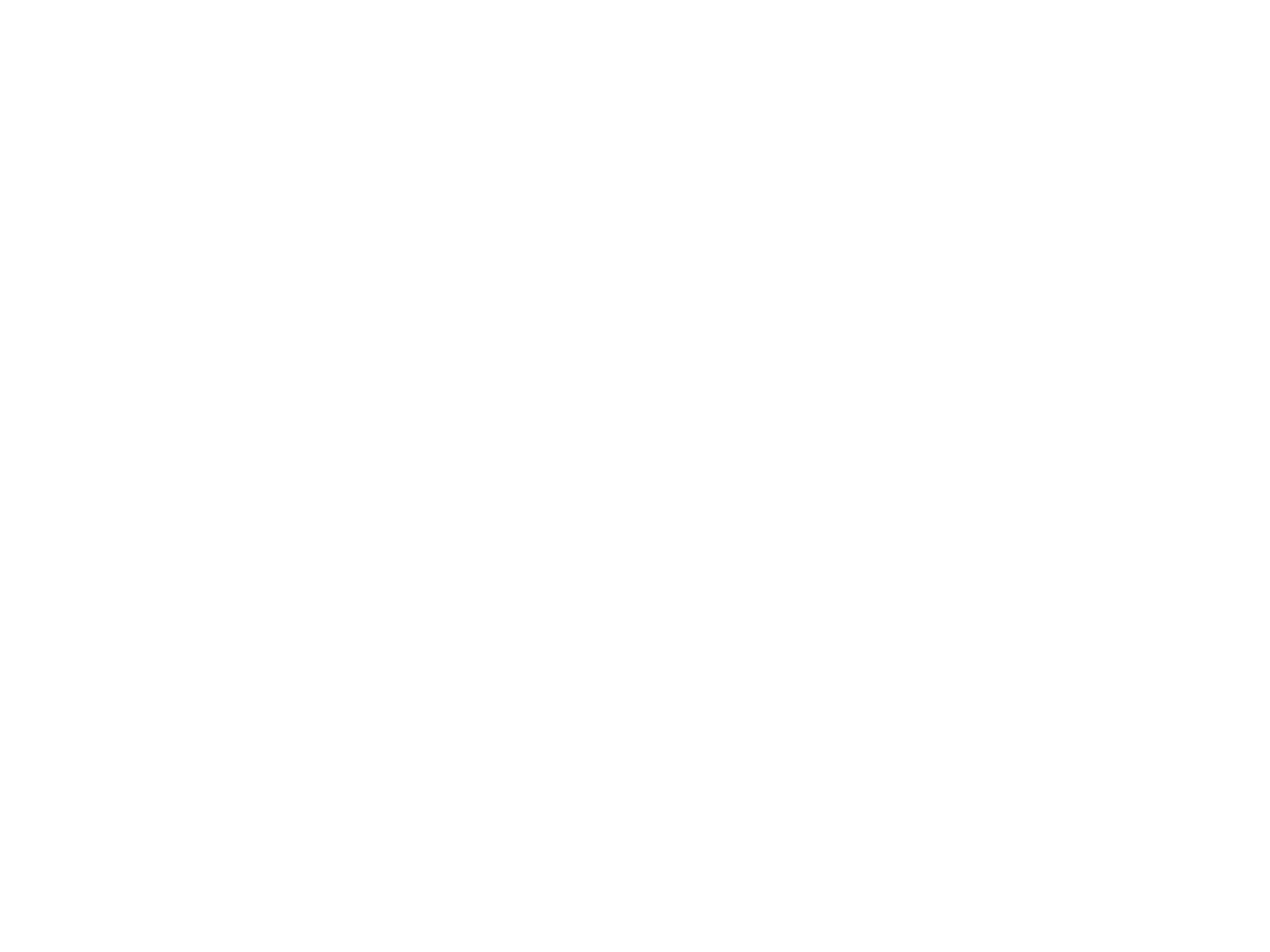

#

### Slide 2
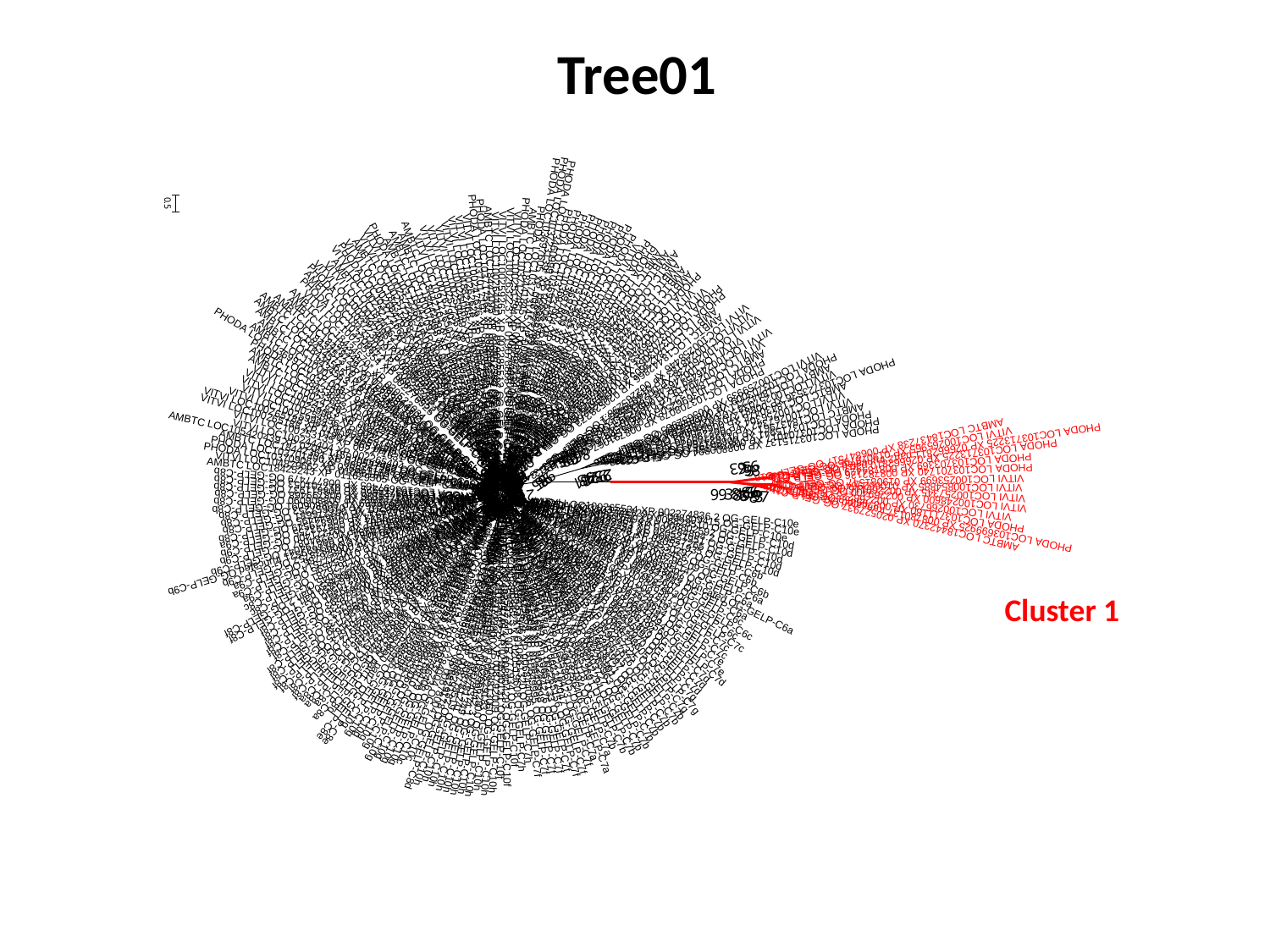

Tree01
Cluster 1

### Slide 3
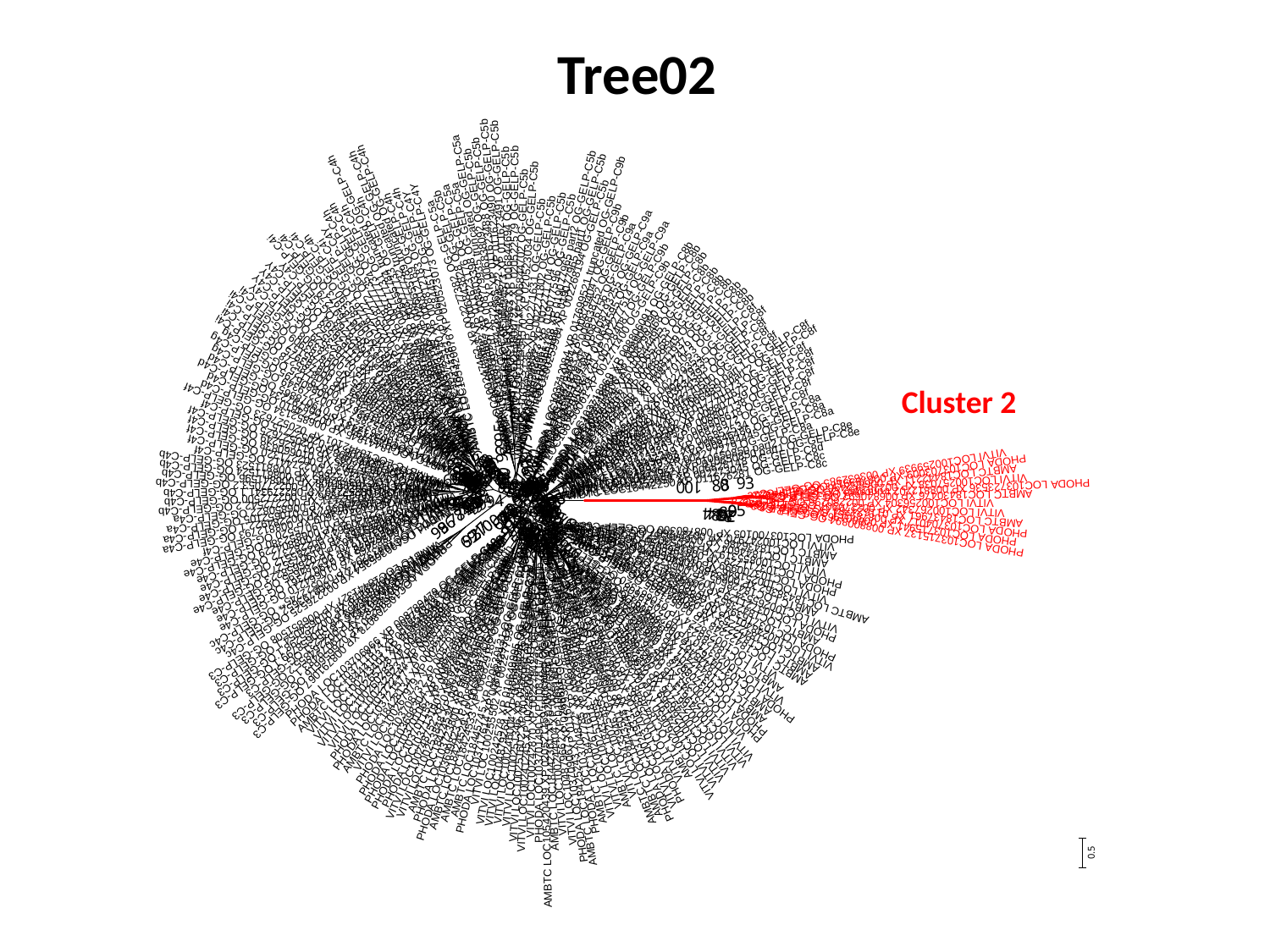

Tree02
Cluster 2

### Slide 4
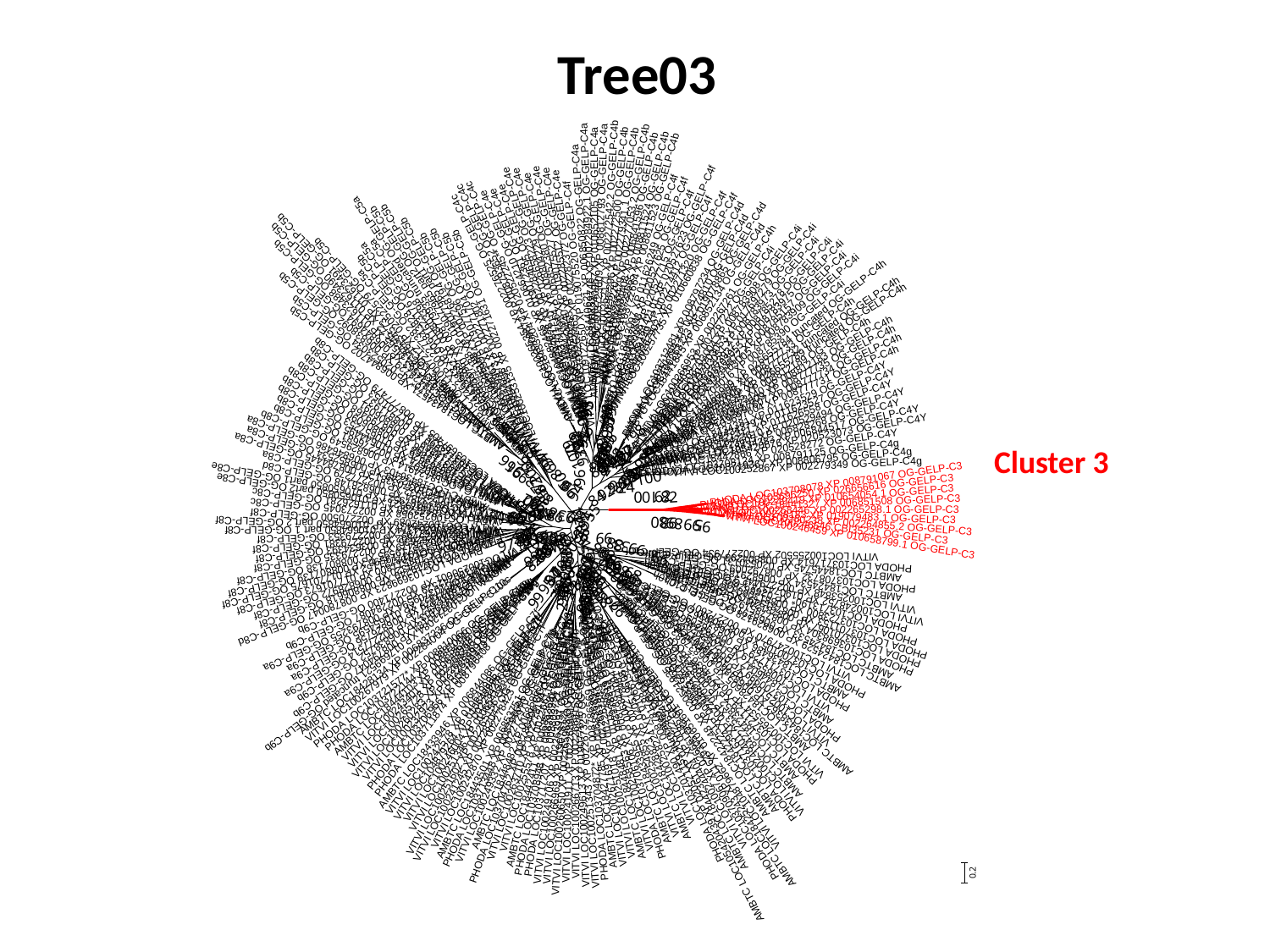

Tree03
Cluster 3

### Slide 5
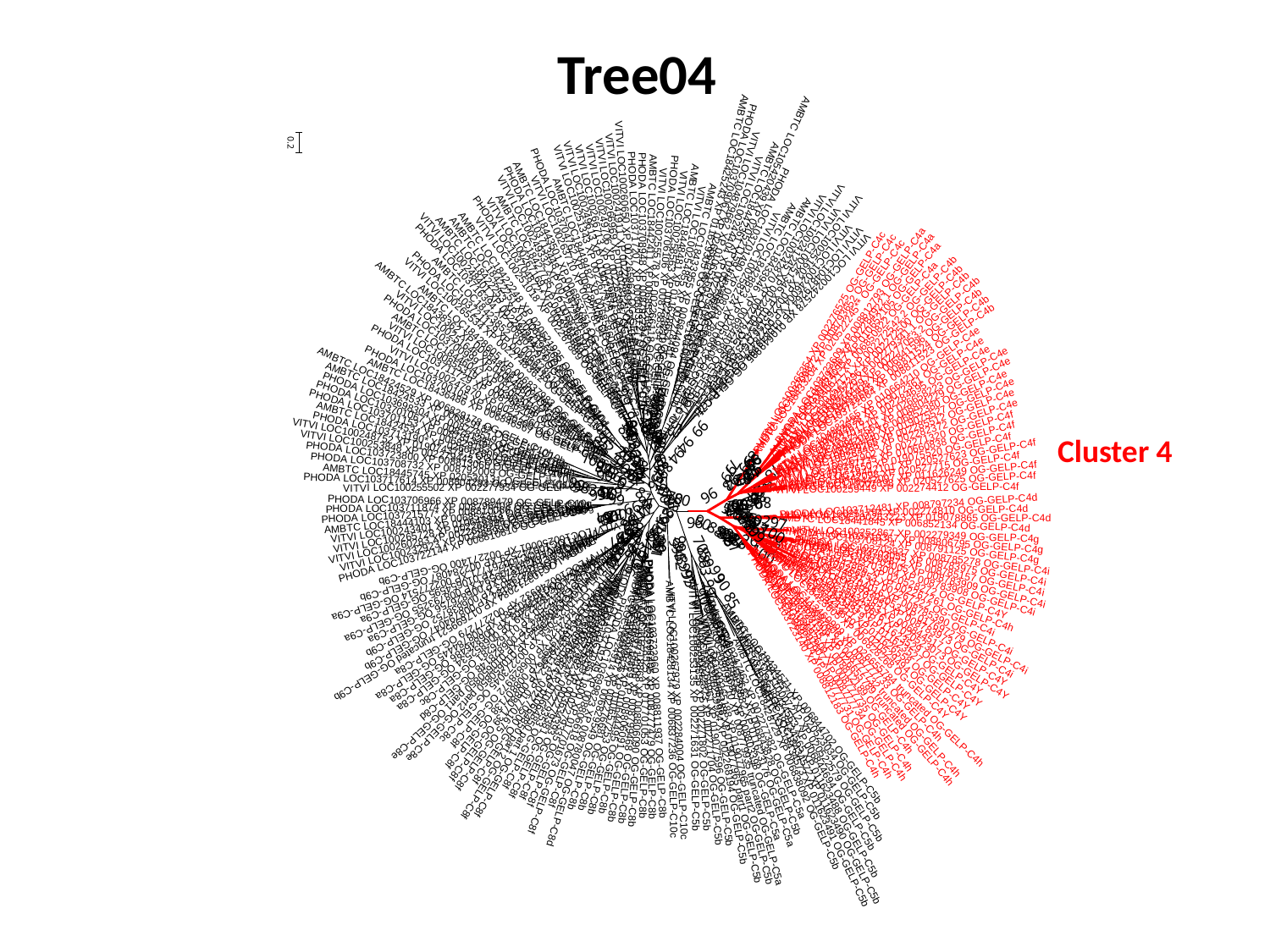

Tree04
Cluster 4

### Slide 6
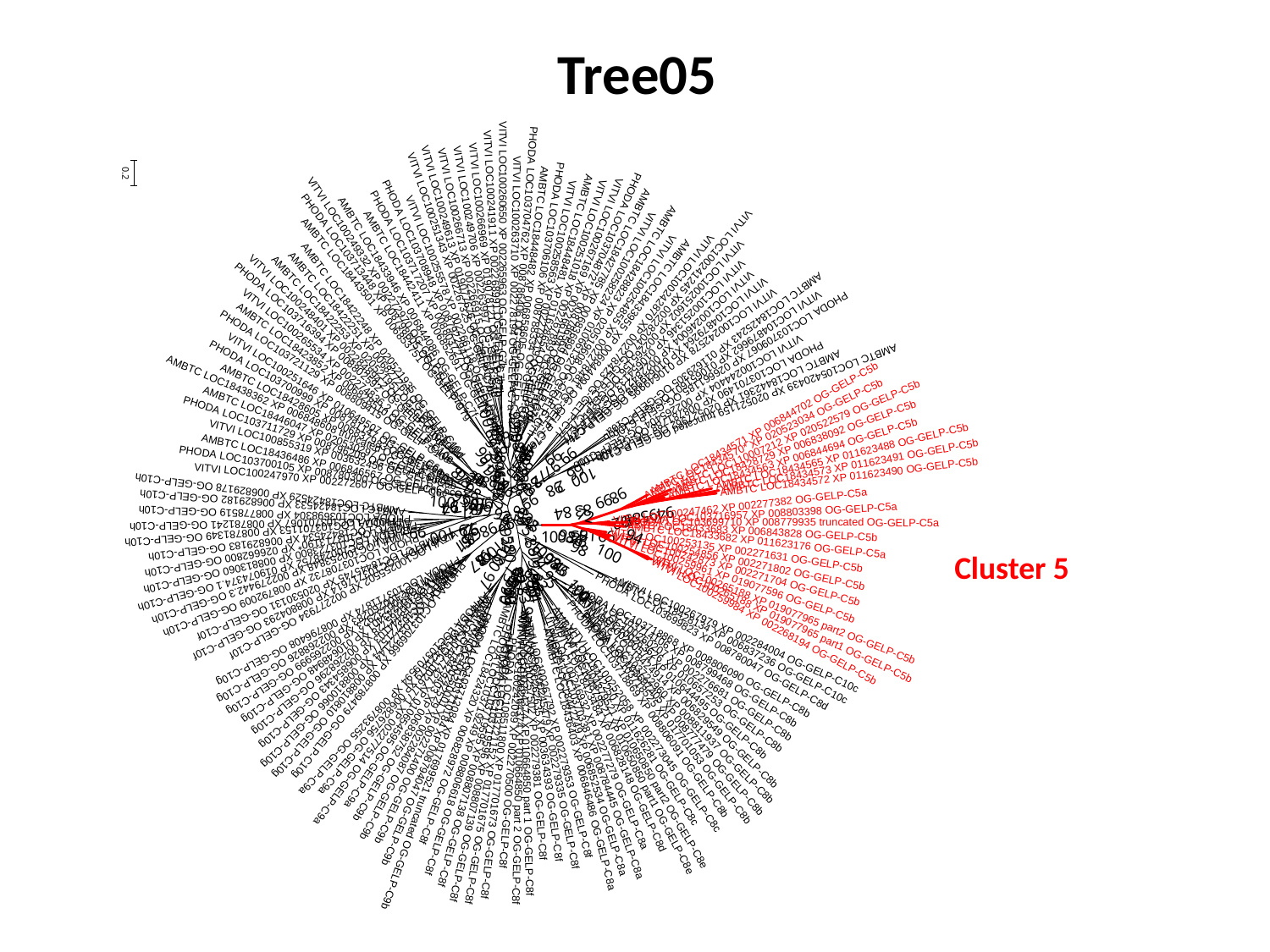

Tree05
Cluster 5

### Slide 7
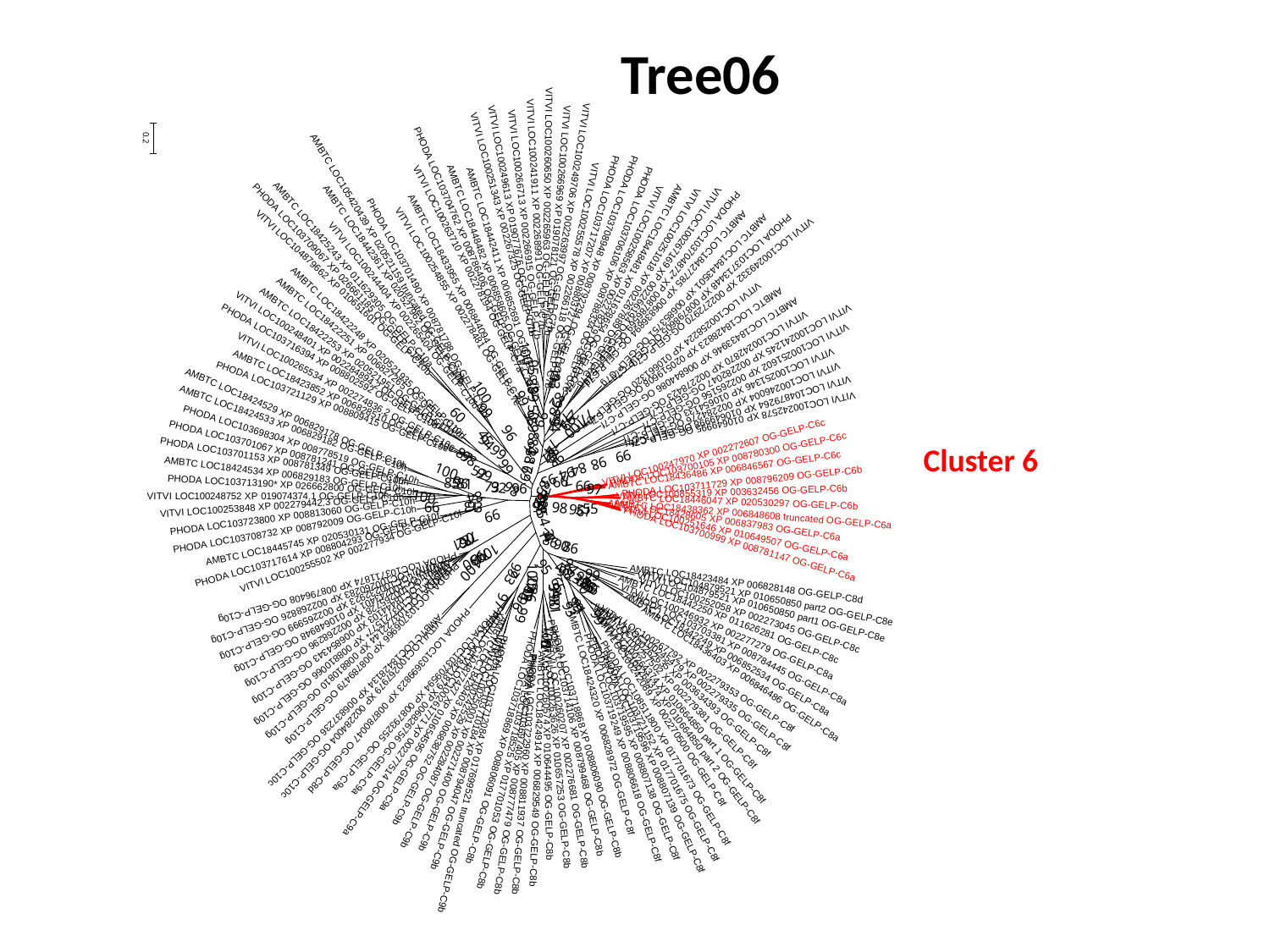

Tree06
Cluster 6

### Slide 8
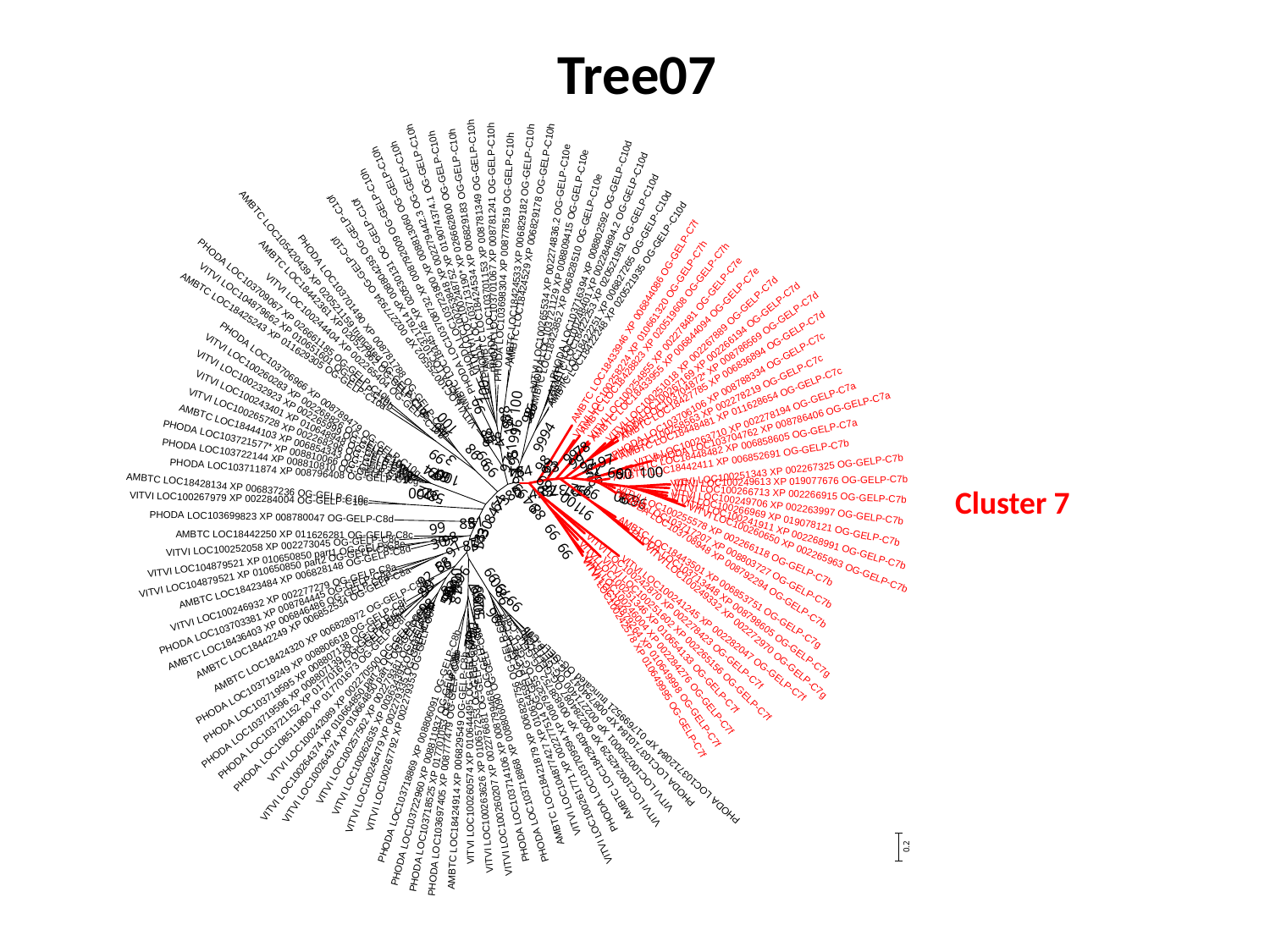

Tree07
Cluster 7

### Slide 9
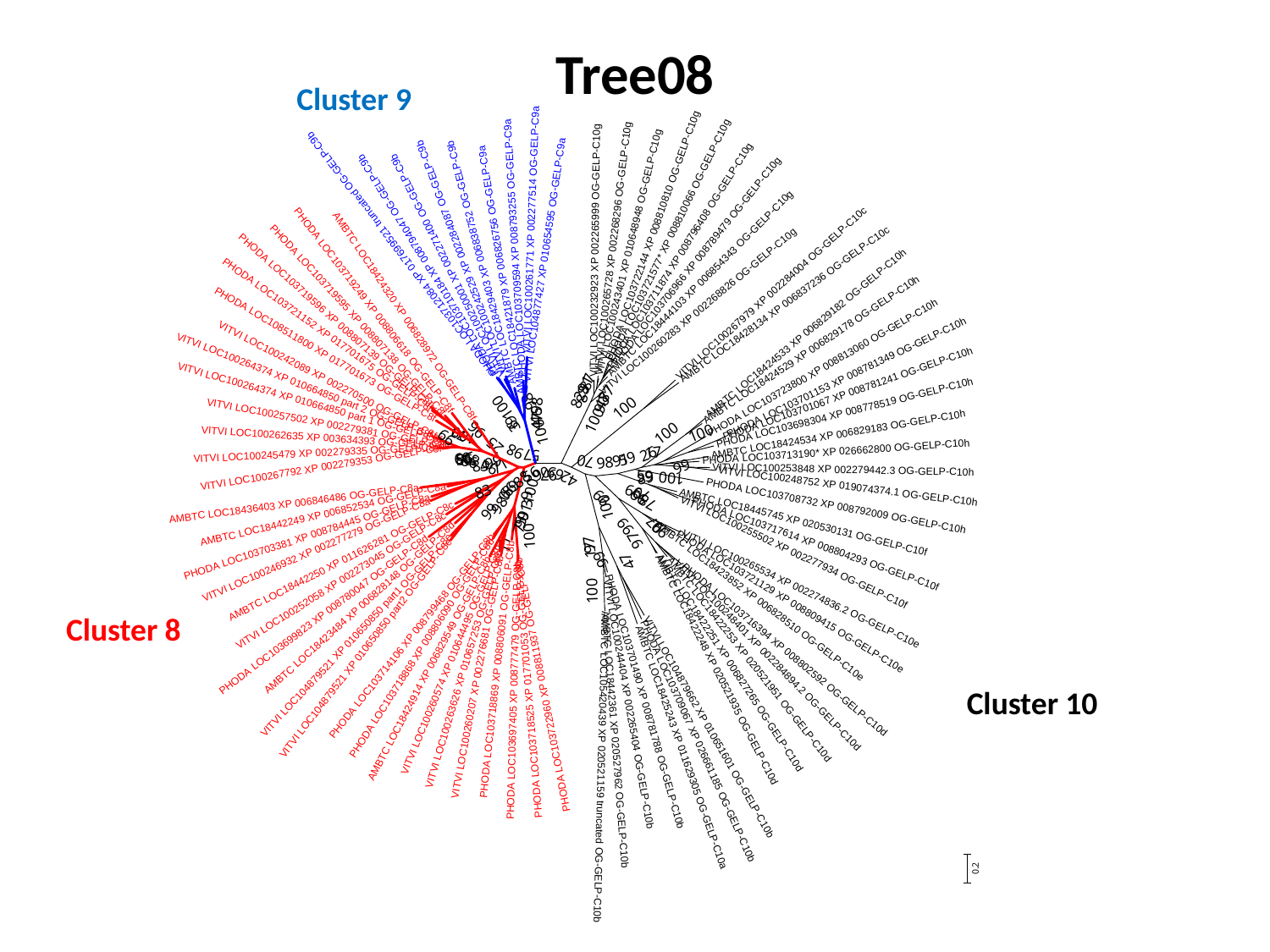

Tree08
Cluster 9
Cluster 8
Cluster 10

### Slide 10
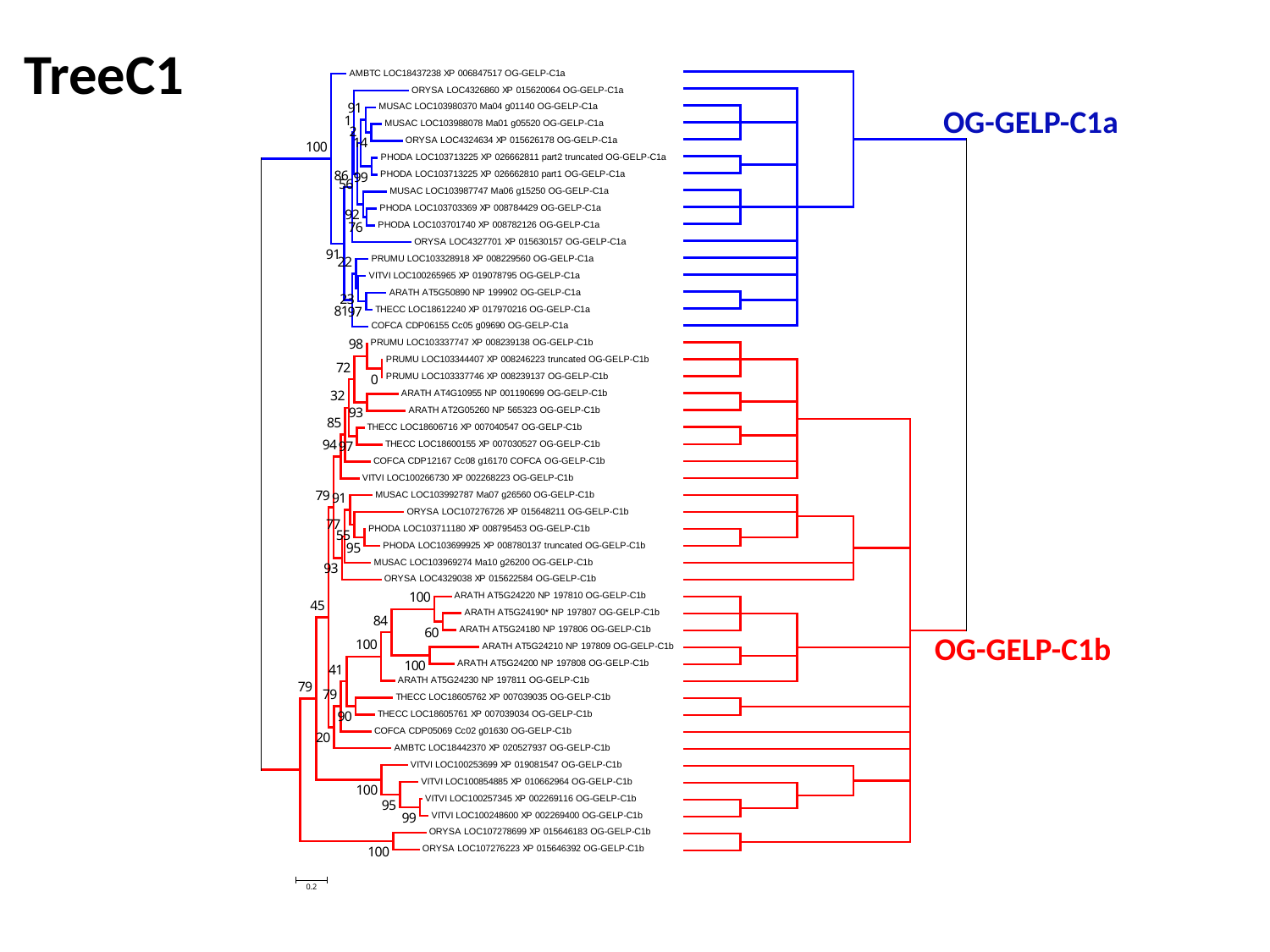

TreeC1
OG-GELP-C1a
OG-GELP-C1b

### Slide 11
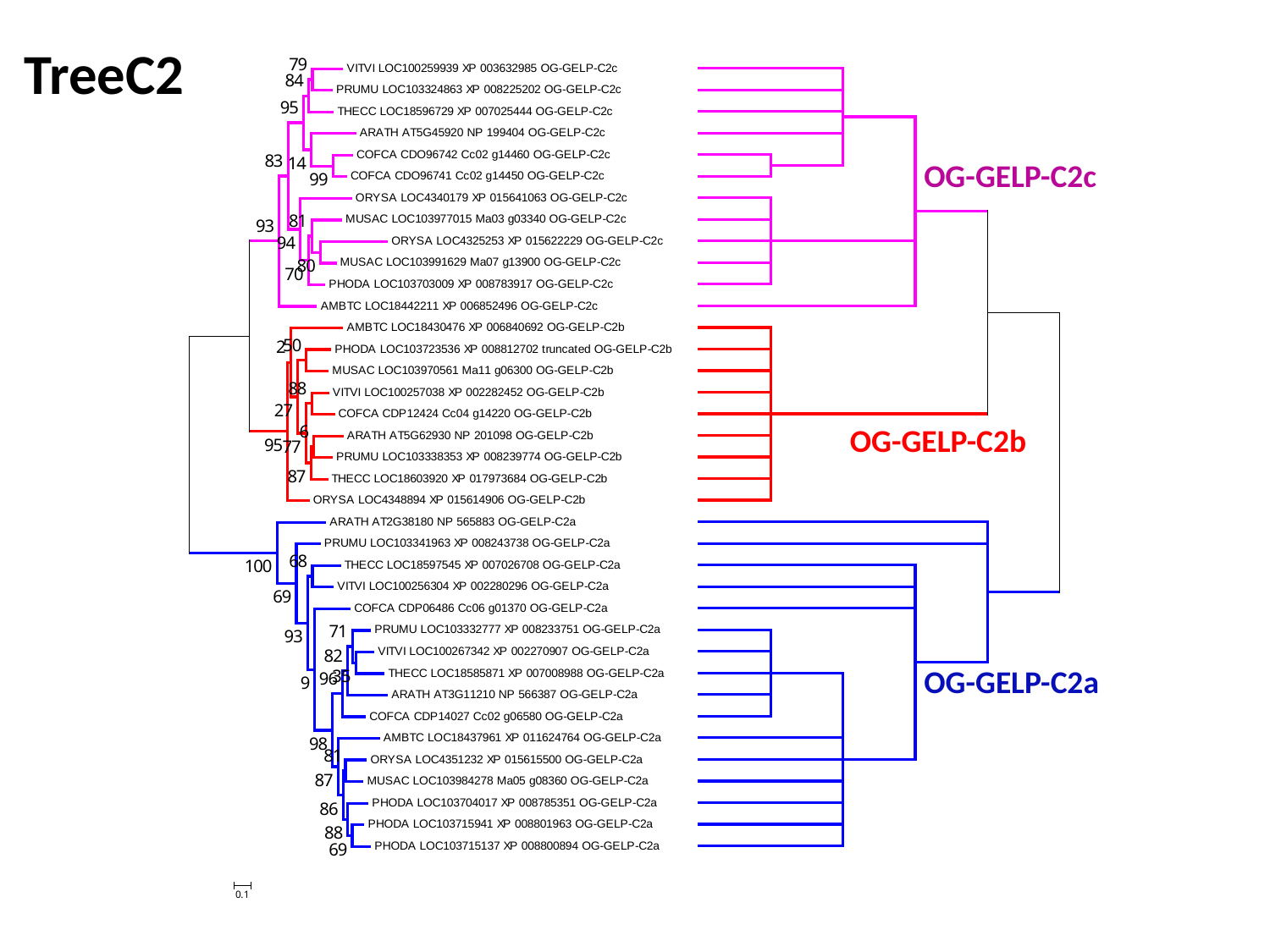

TreeC2
OG-GELP-C2c
OG-GELP-C2b
OG-GELP-C2a

### Slide 12
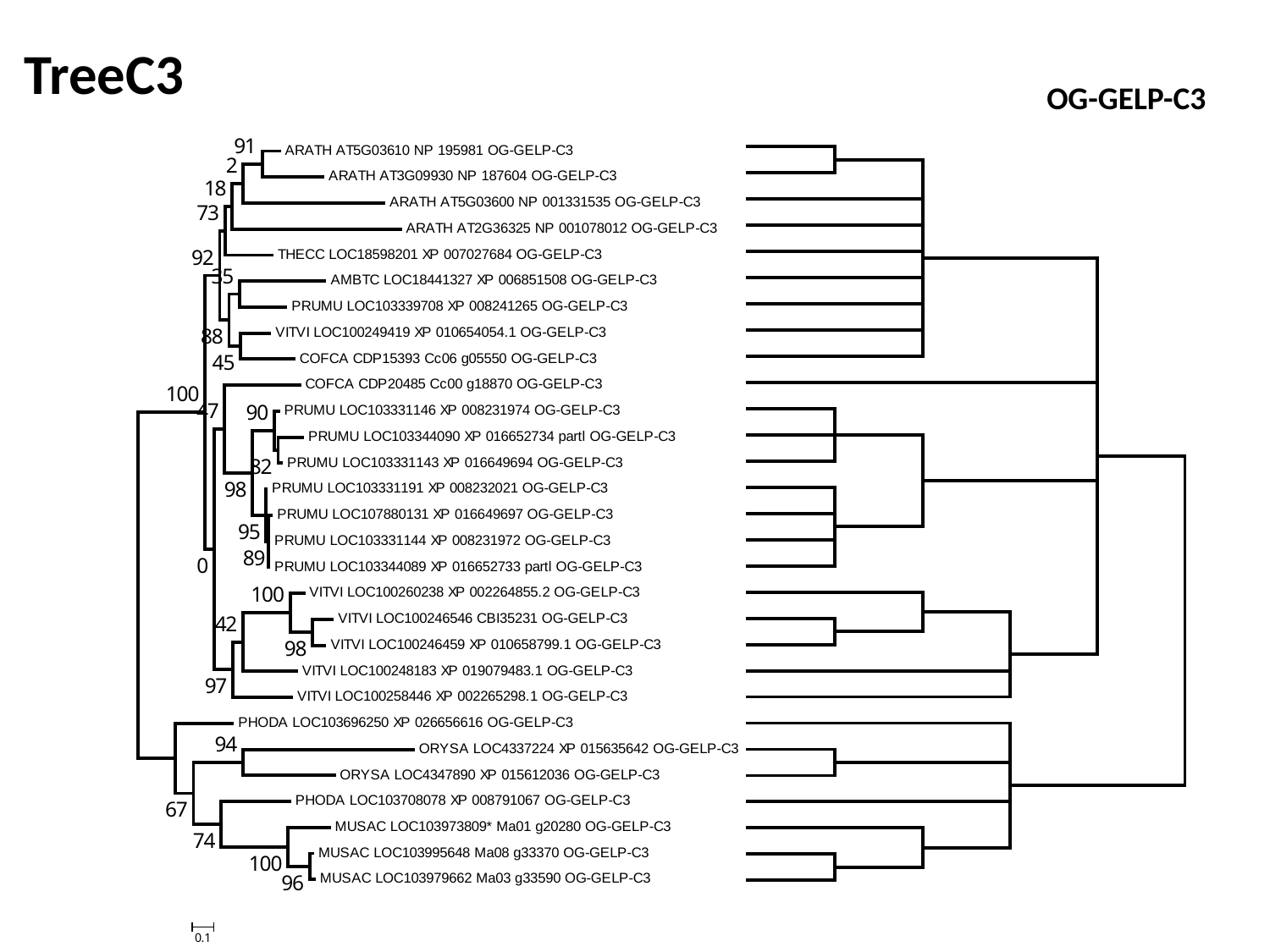

TreeC3
OG-GELP-C3

### Slide 13
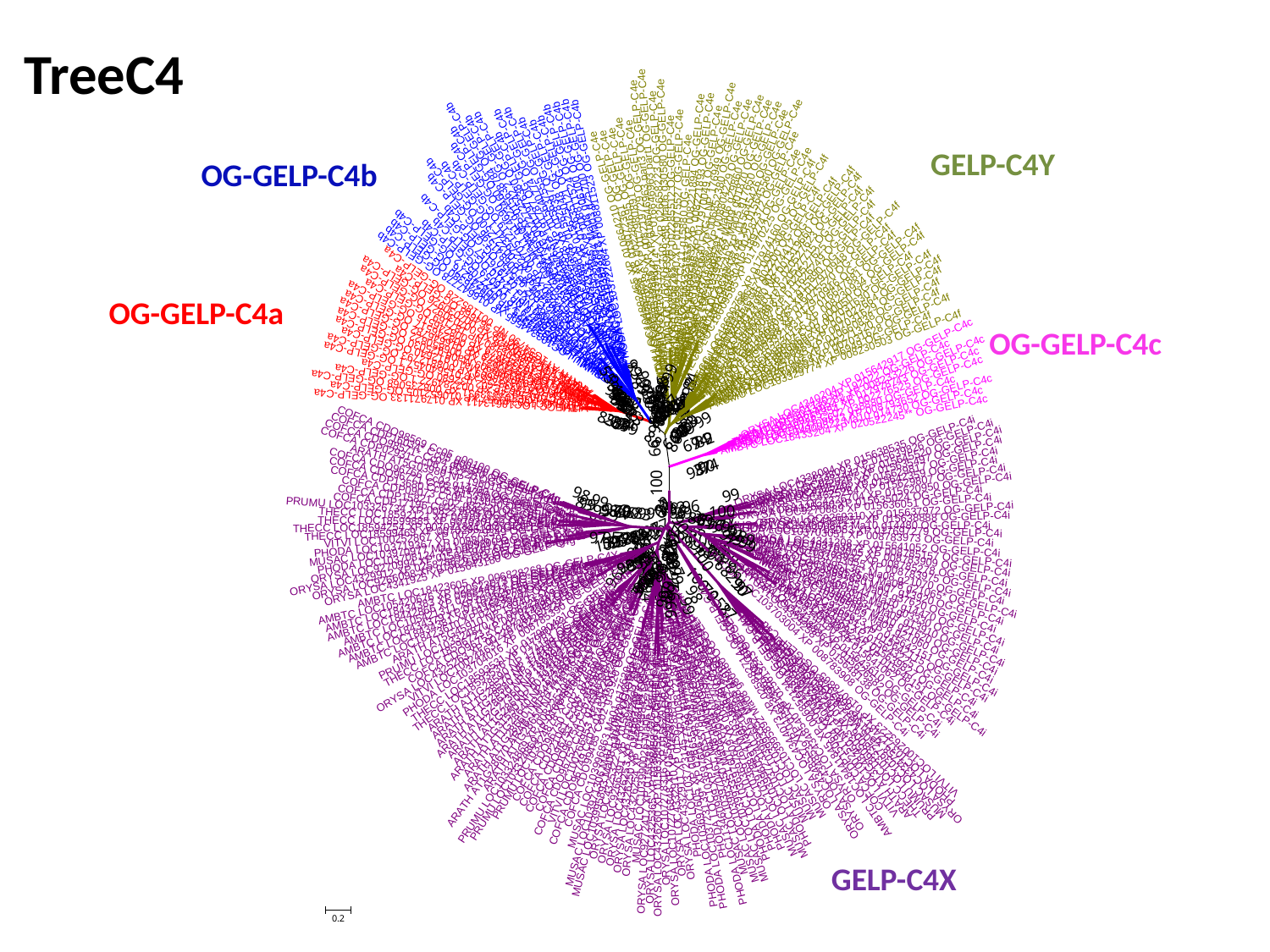

TreeC4
GELP-C4Y
OG-GELP-C4b
OG-GELP-C4a
OG-GELP-C4c
GELP-C4X

### Slide 14
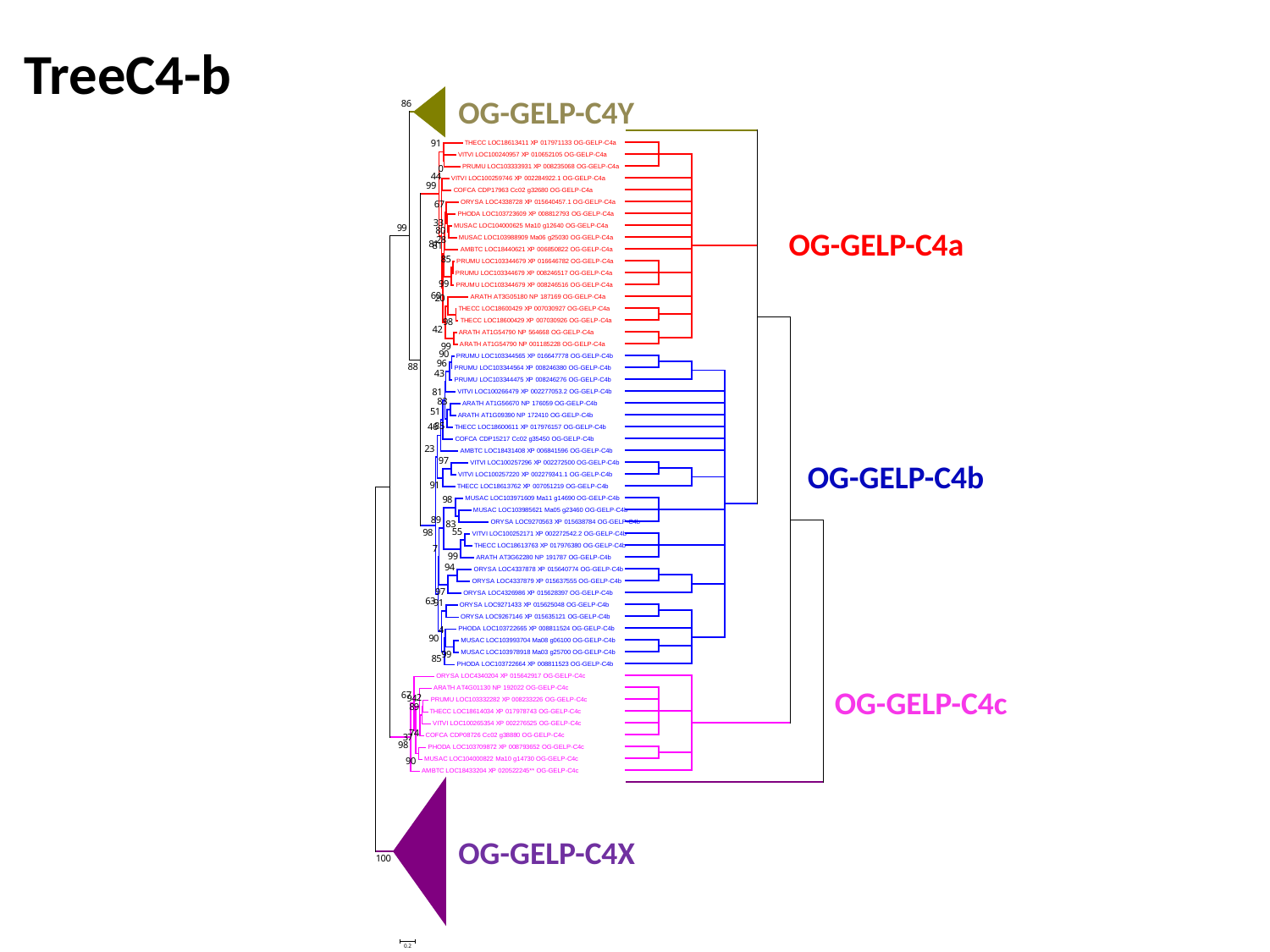

TreeC4-b
OG-GELP-C4Y
OG-GELP-C4a
OG-GELP-C4b
OG-GELP-C4c
OG-GELP-C4X

### Slide 15
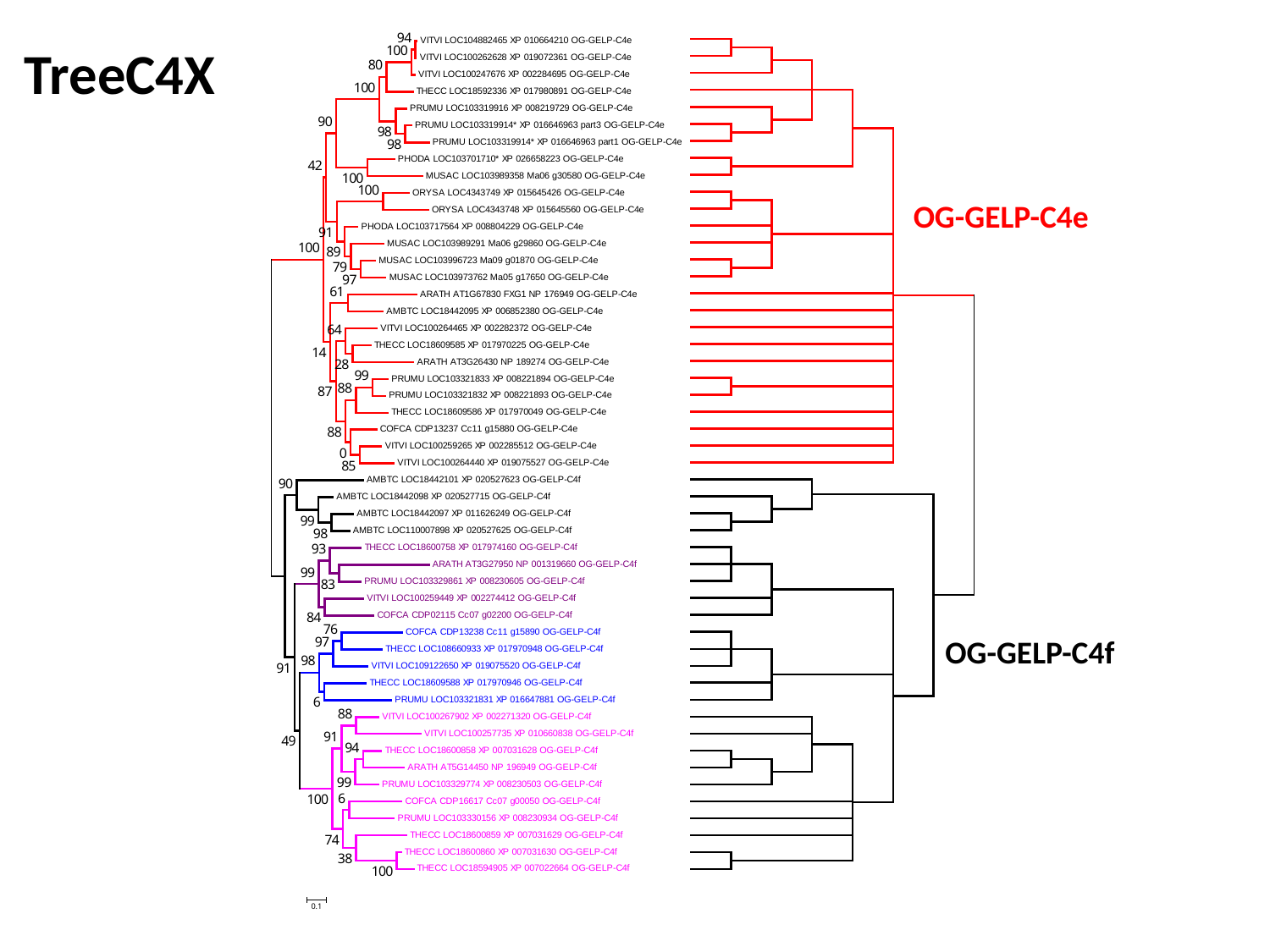

TreeC4X
OG-GELP-C4e
OG-GELP-C4f

### Slide 16
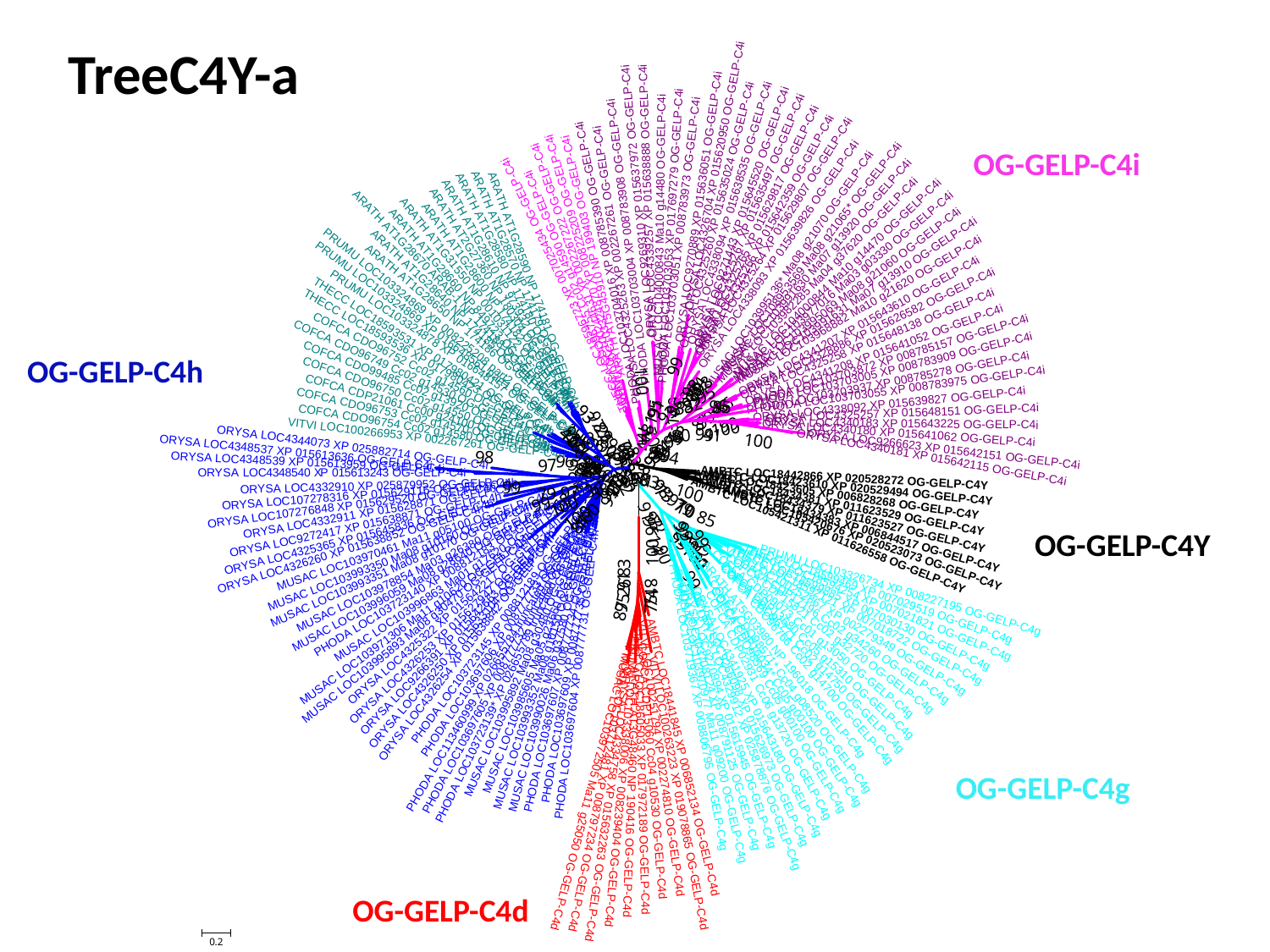

TreeC4Y-a
OG-GELP-C4i
OG-GELP-C4h
OG-GELP-C4Y
OG-GELP-C4g
OG-GELP-C4d

### Slide 17
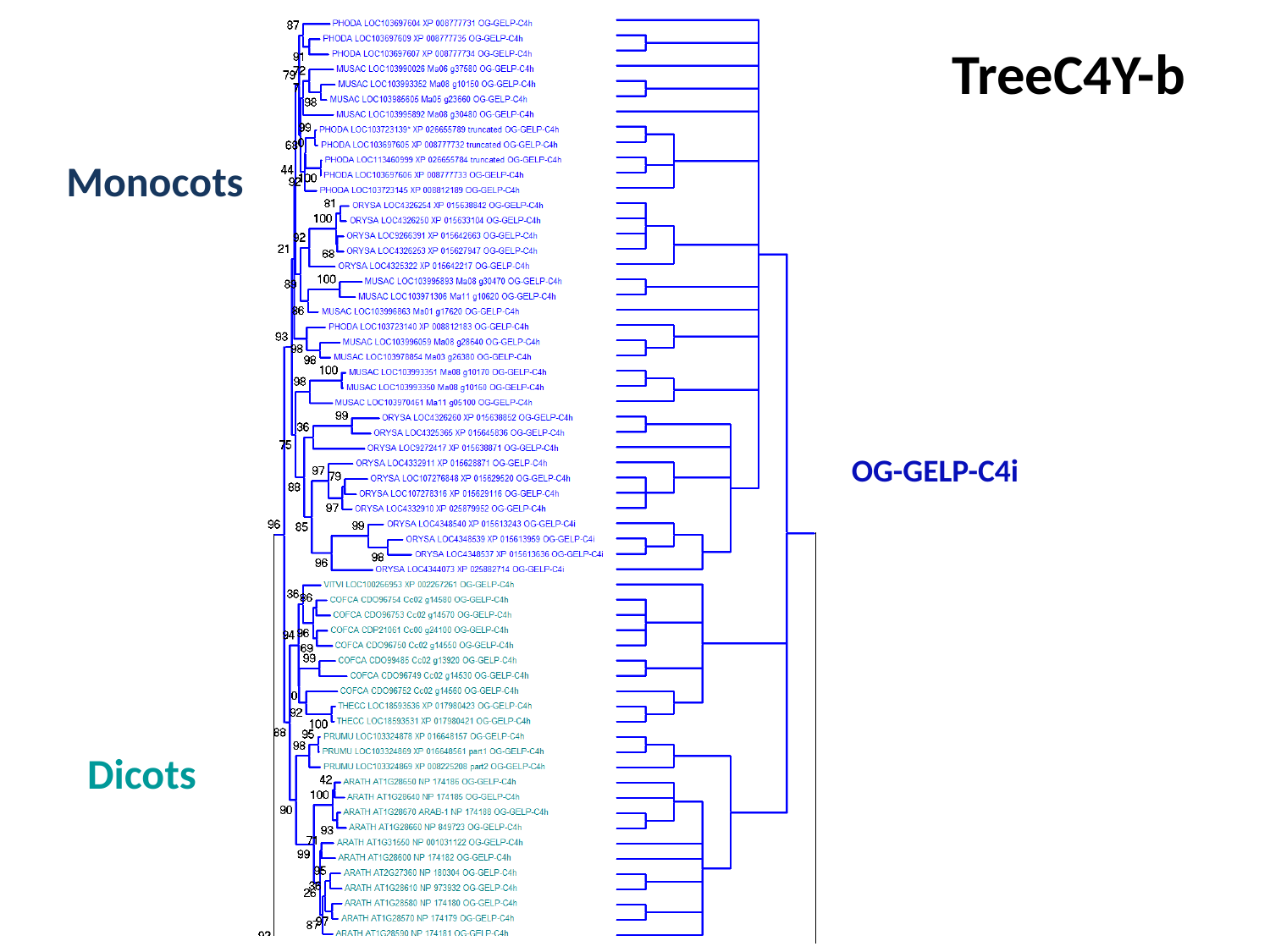

TreeC4Y-b
Monocots
OG-GELP-C4i
Dicots

### Slide 18
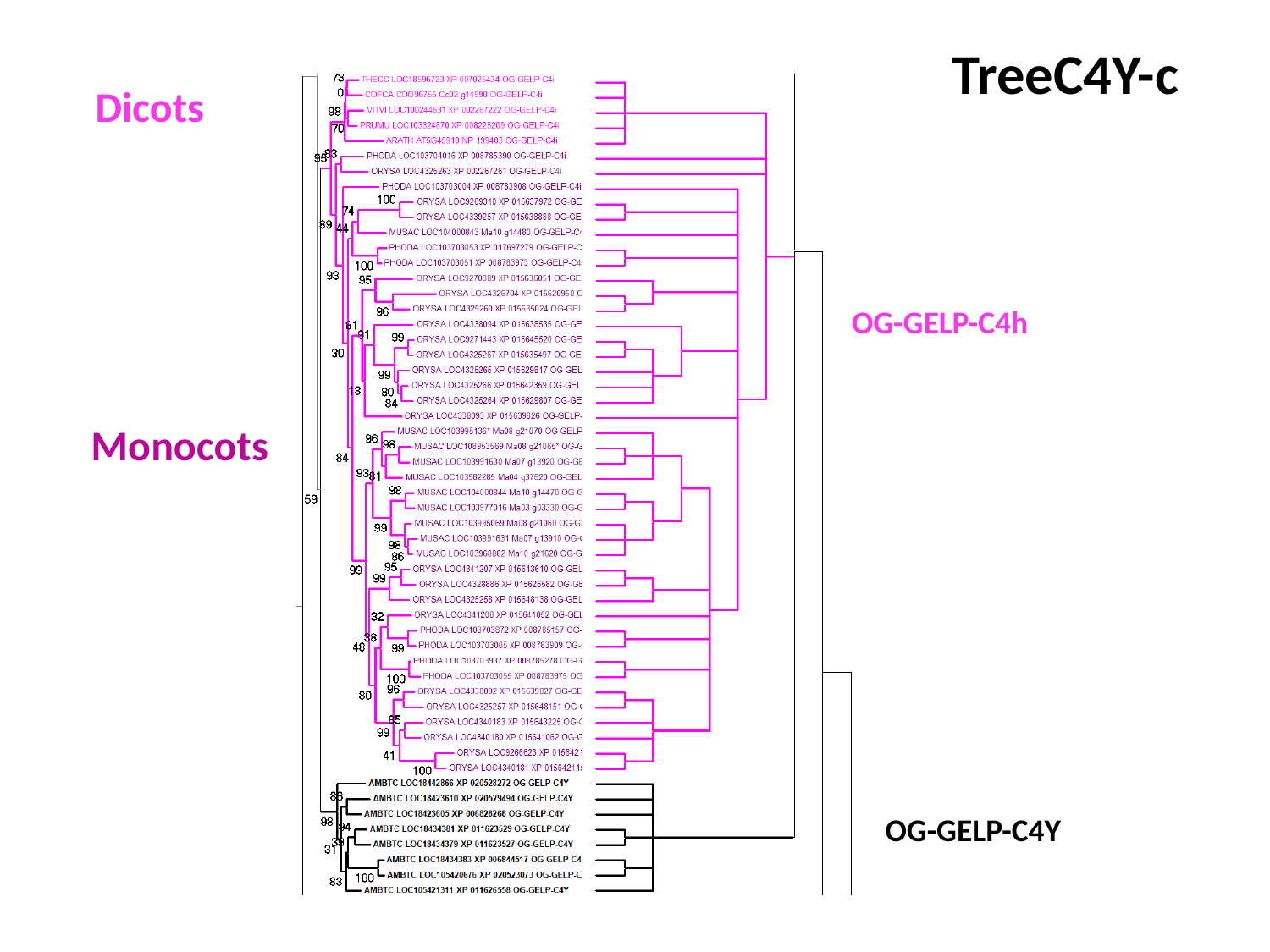

TreeC4Y-c
Dicots
OG-GELP-C4h
Monocots
OG-GELP-C4Y

### Slide 19
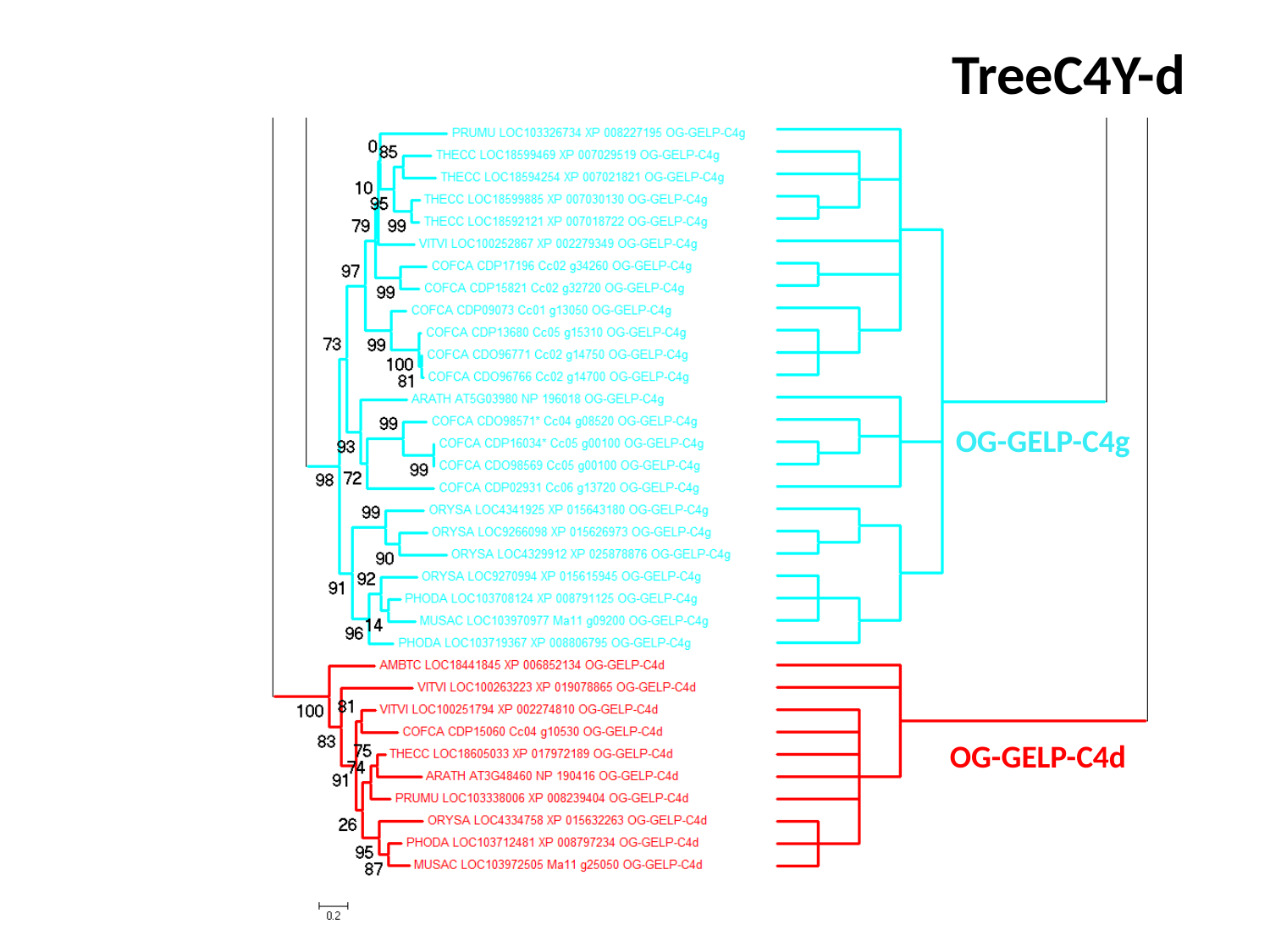

TreeC4Y-d
OG-GELP-C4g
OG-GELP-C4d

### Slide 20
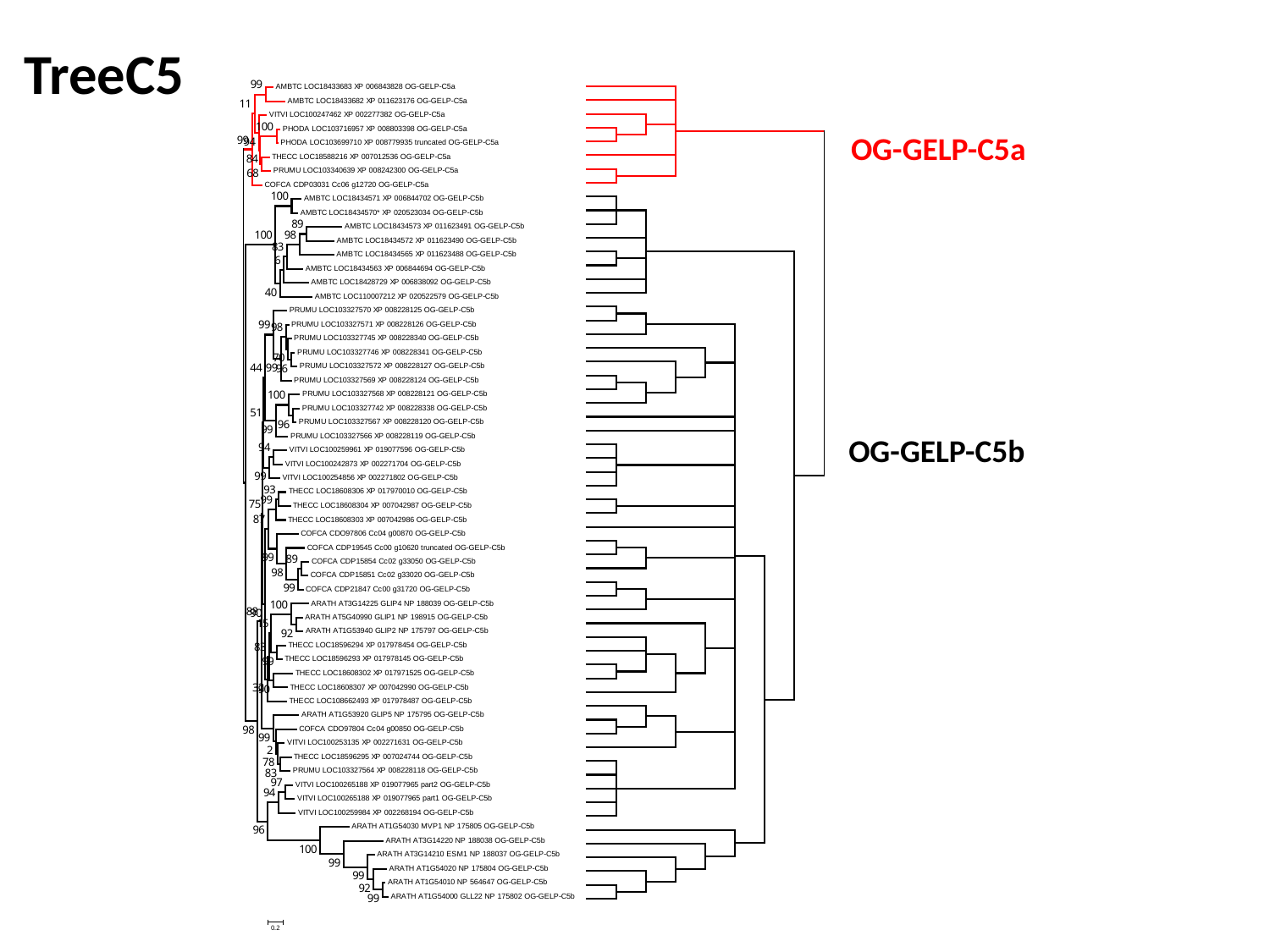

TreeC5
OG-GELP-C5a
OG-GELP-C5b

### Slide 21
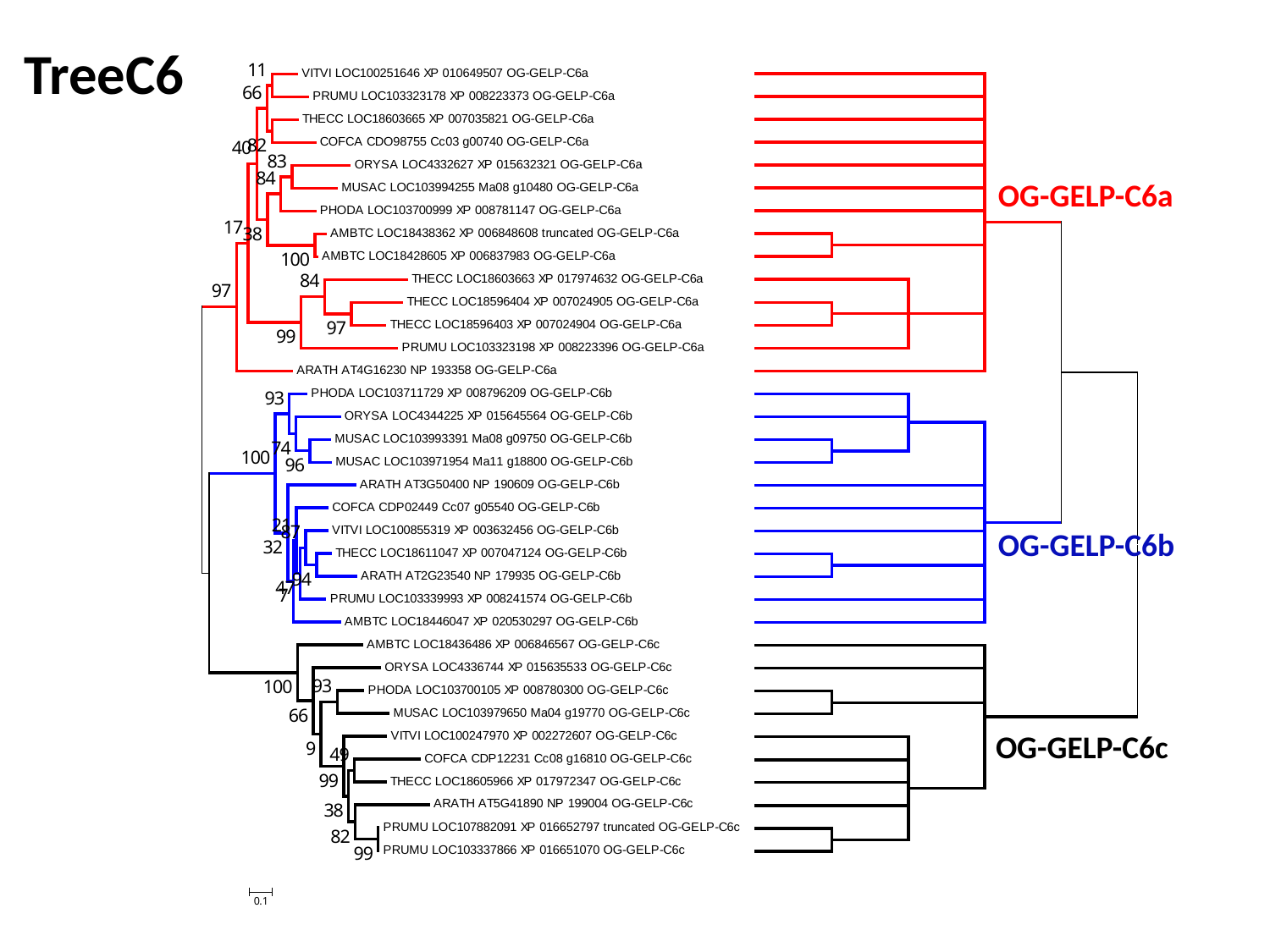

TreeC6
OG-GELP-C6a
OG-GELP-C6b
OG-GELP-C6c

### Slide 22
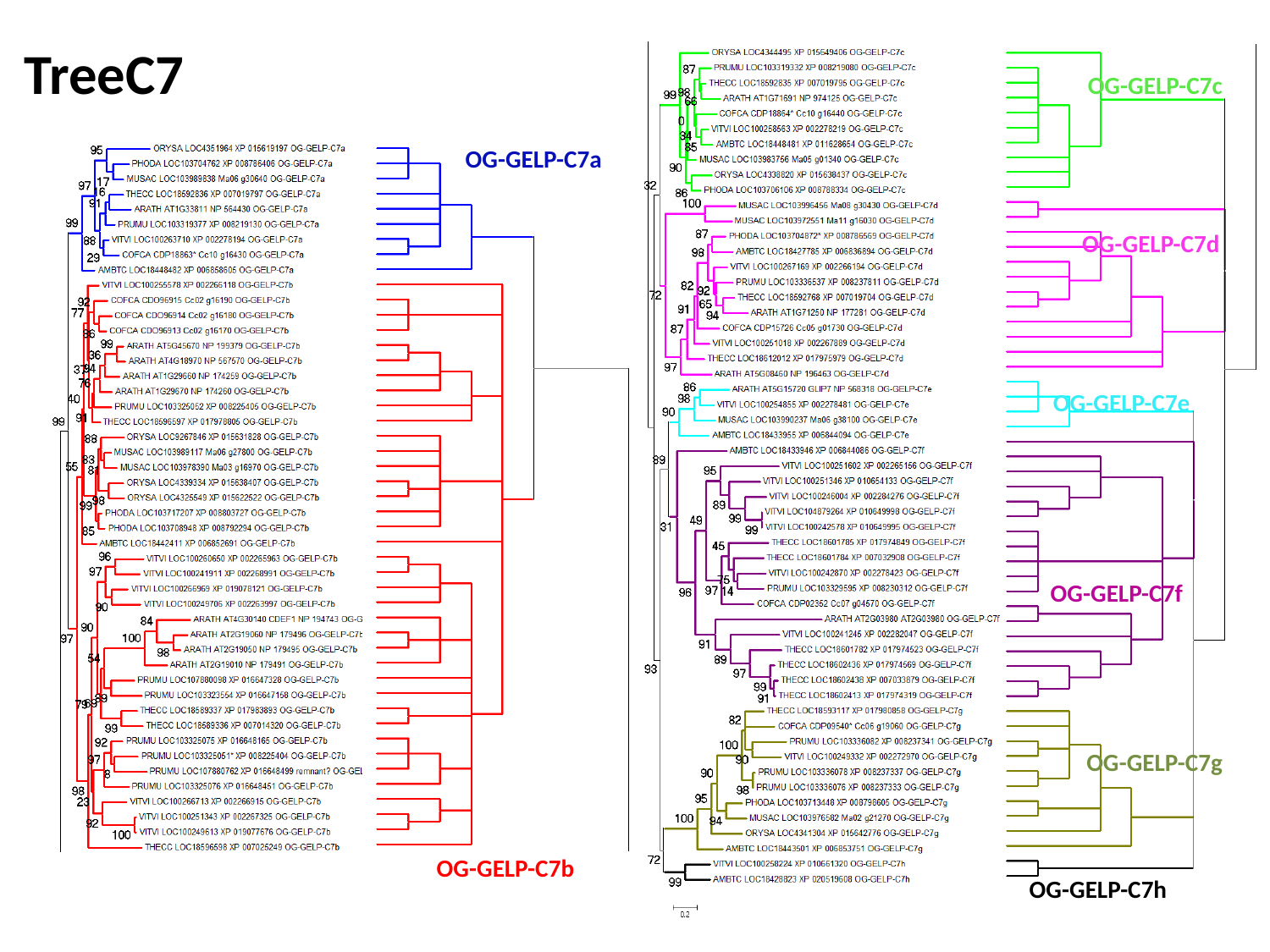

TreeC7
OG-GELP-C7c
OG-GELP-C7a
OG-GELP-C7d
OG-GELP-C7e
OG-GELP-C7f
OG-GELP-C7g
OG-GELP-C7b
OG-GELP-C7h

### Slide 23
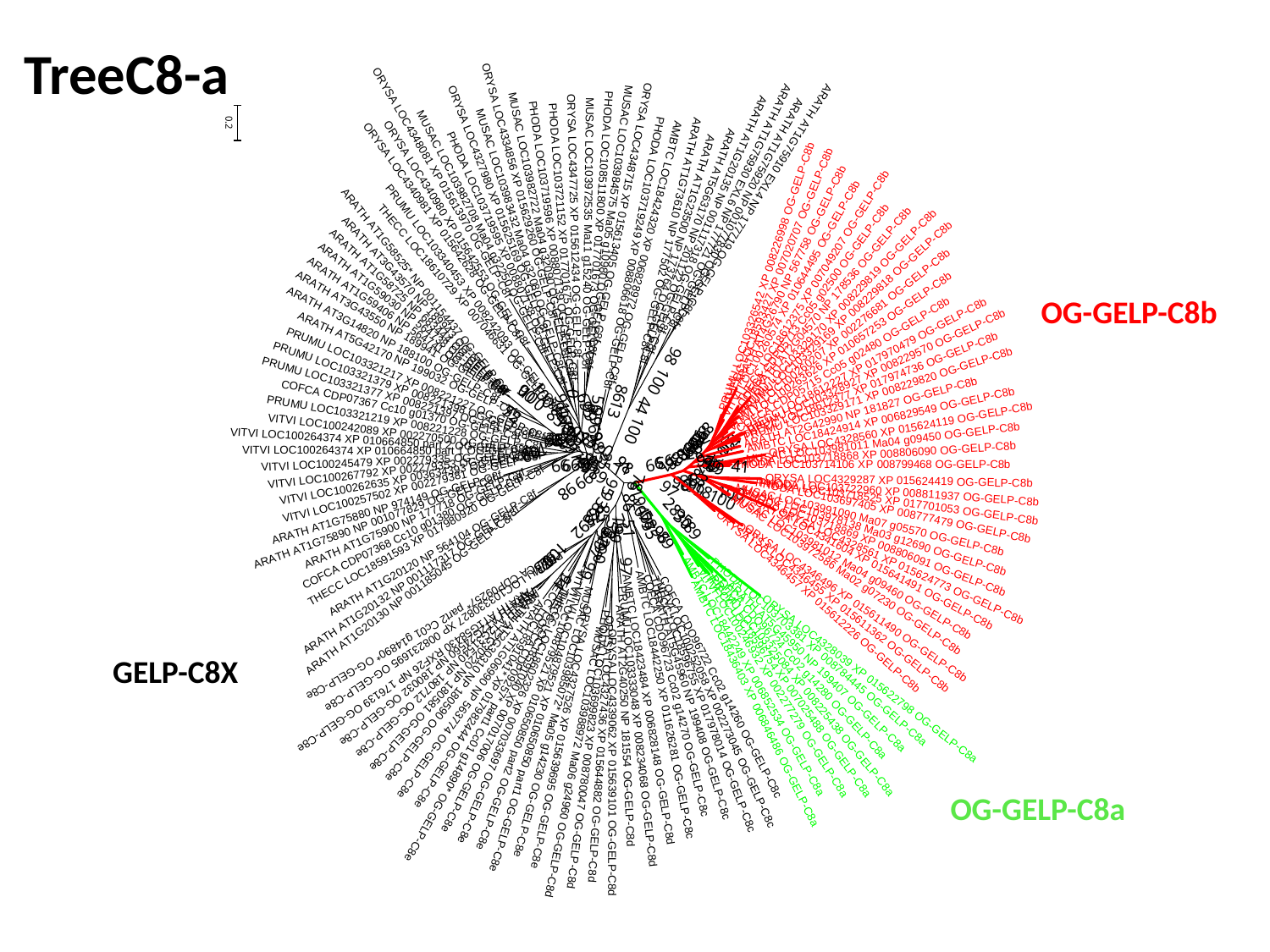

TreeC8-a
OG-GELP-C8b
GELP-C8X
OG-GELP-C8a

### Slide 24
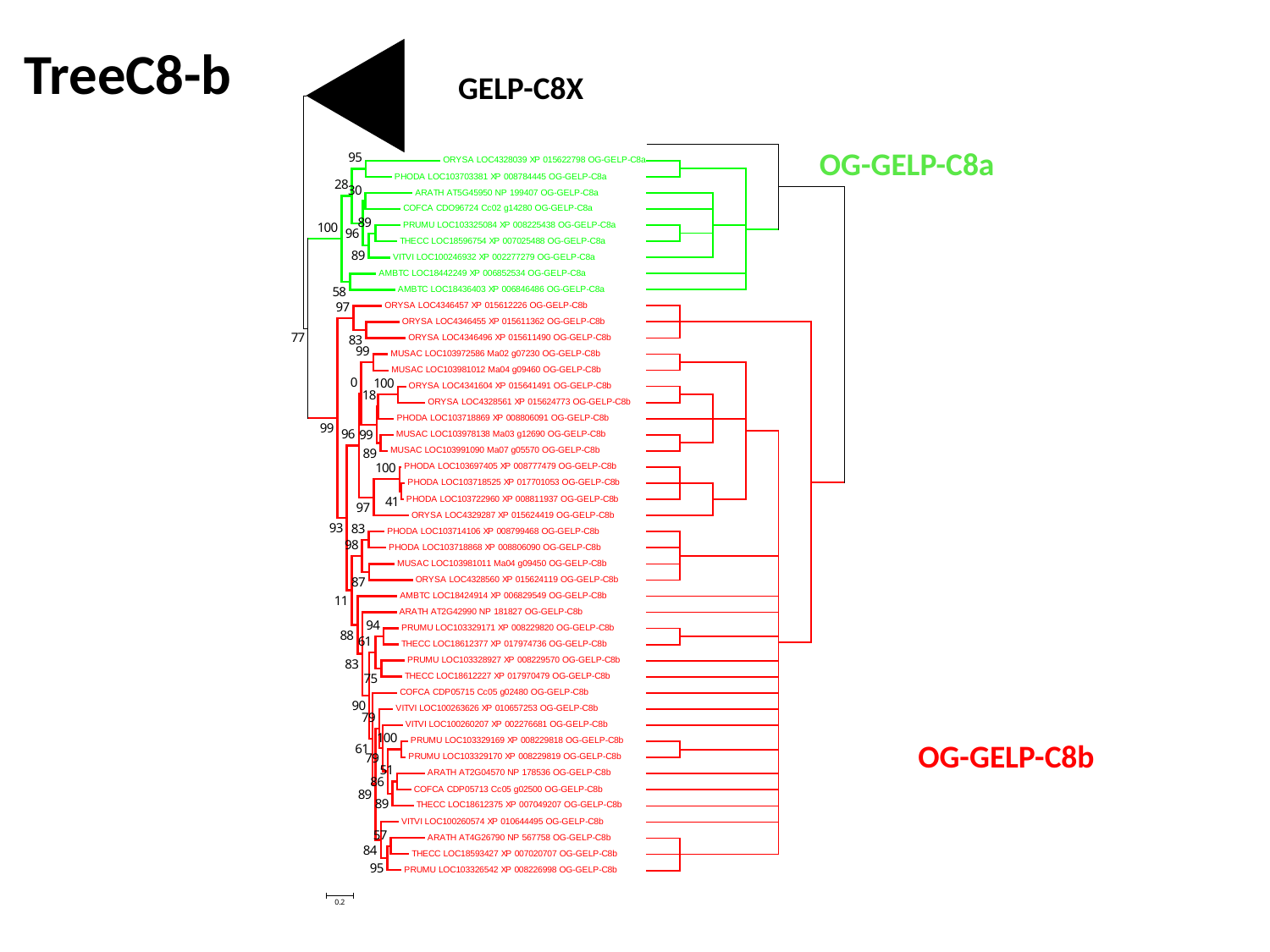

TreeC8-b
GELP-C8X
OG-GELP-C8a
OG-GELP-C8b

### Slide 25
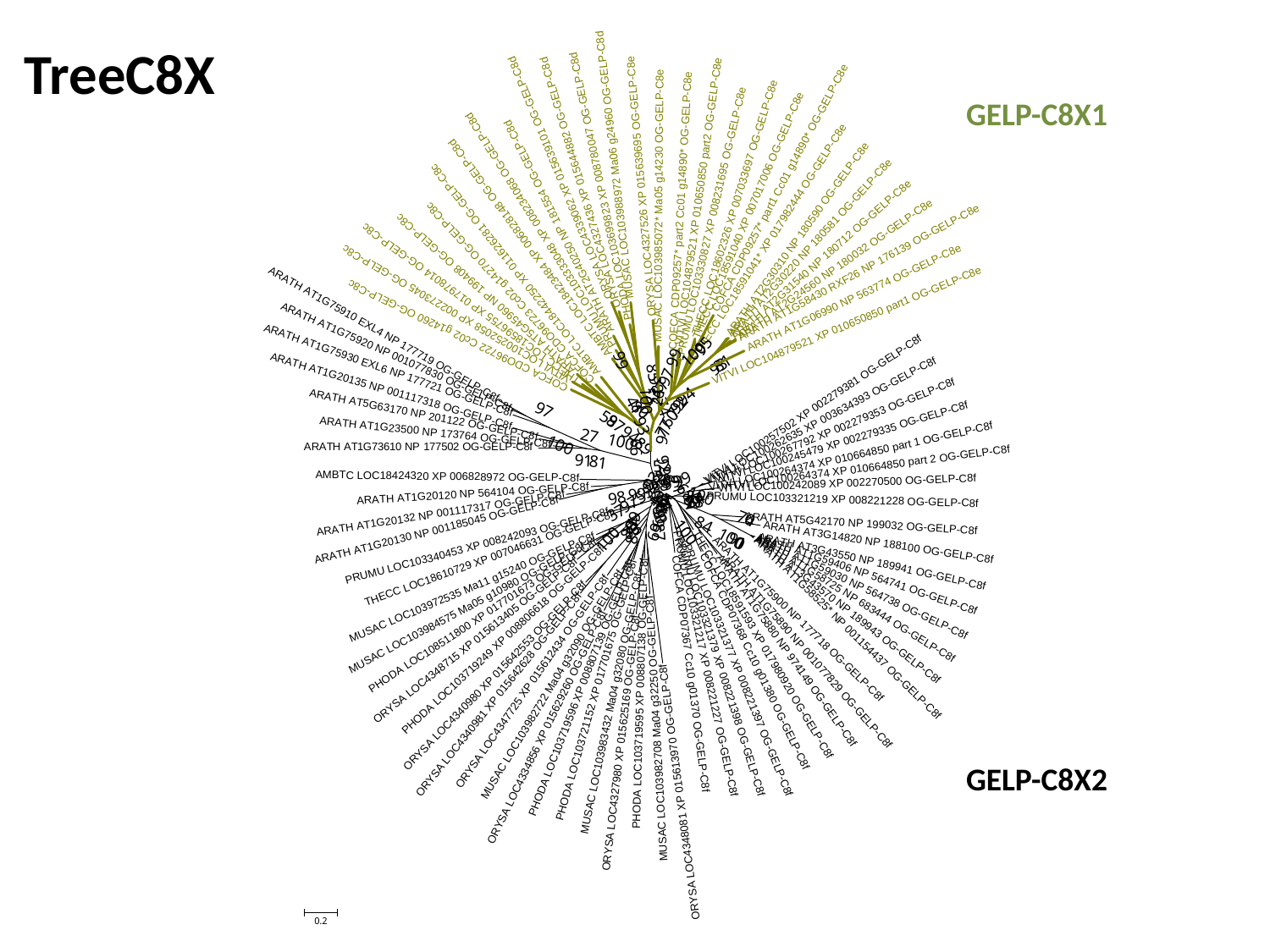

TreeC8X
GELP-C8X1
GELP-C8X2

### Slide 26
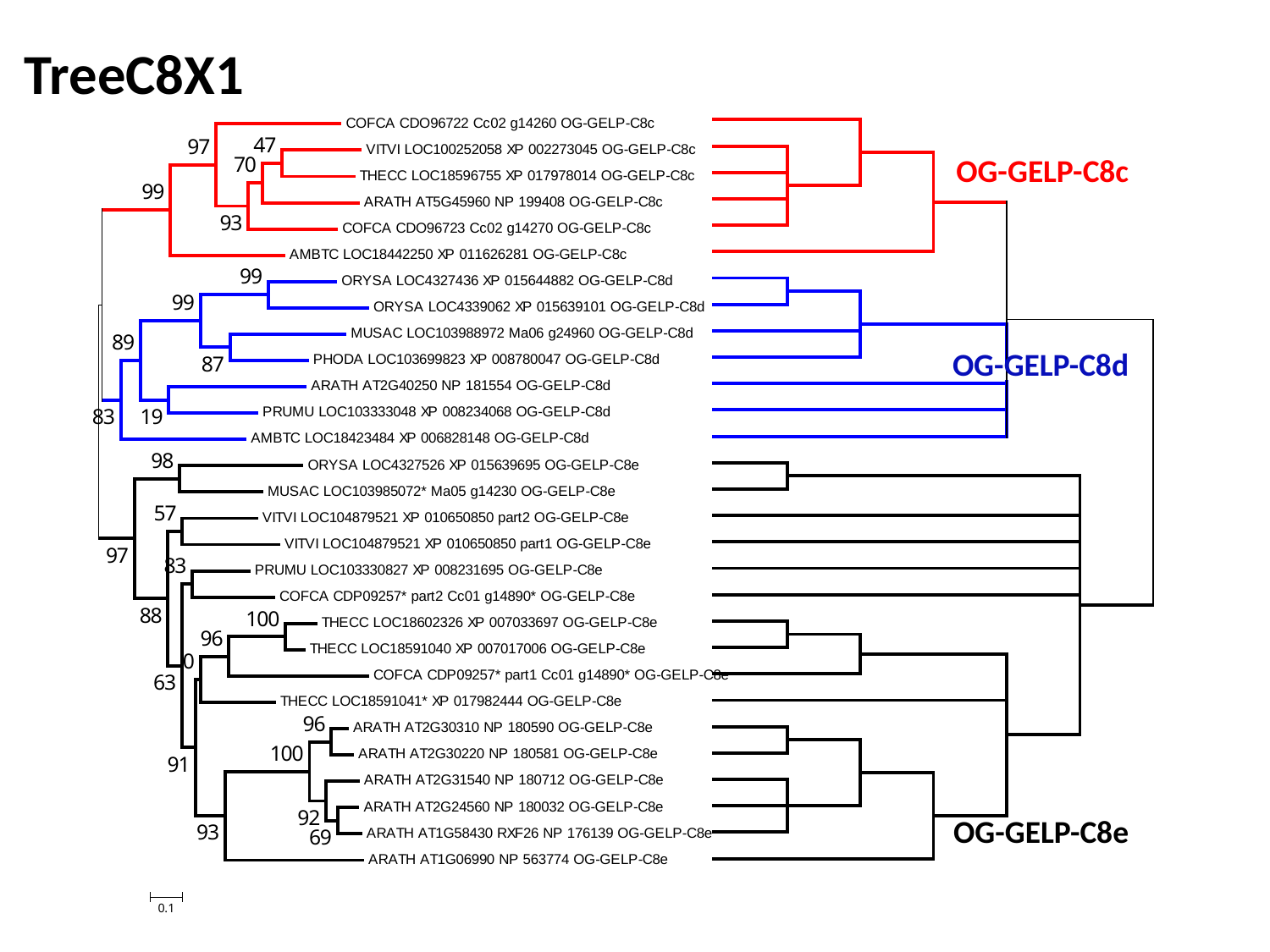

TreeC8X1
OG-GELP-C8c
OG-GELP-C8d
OG-GELP-C8e

### Slide 27
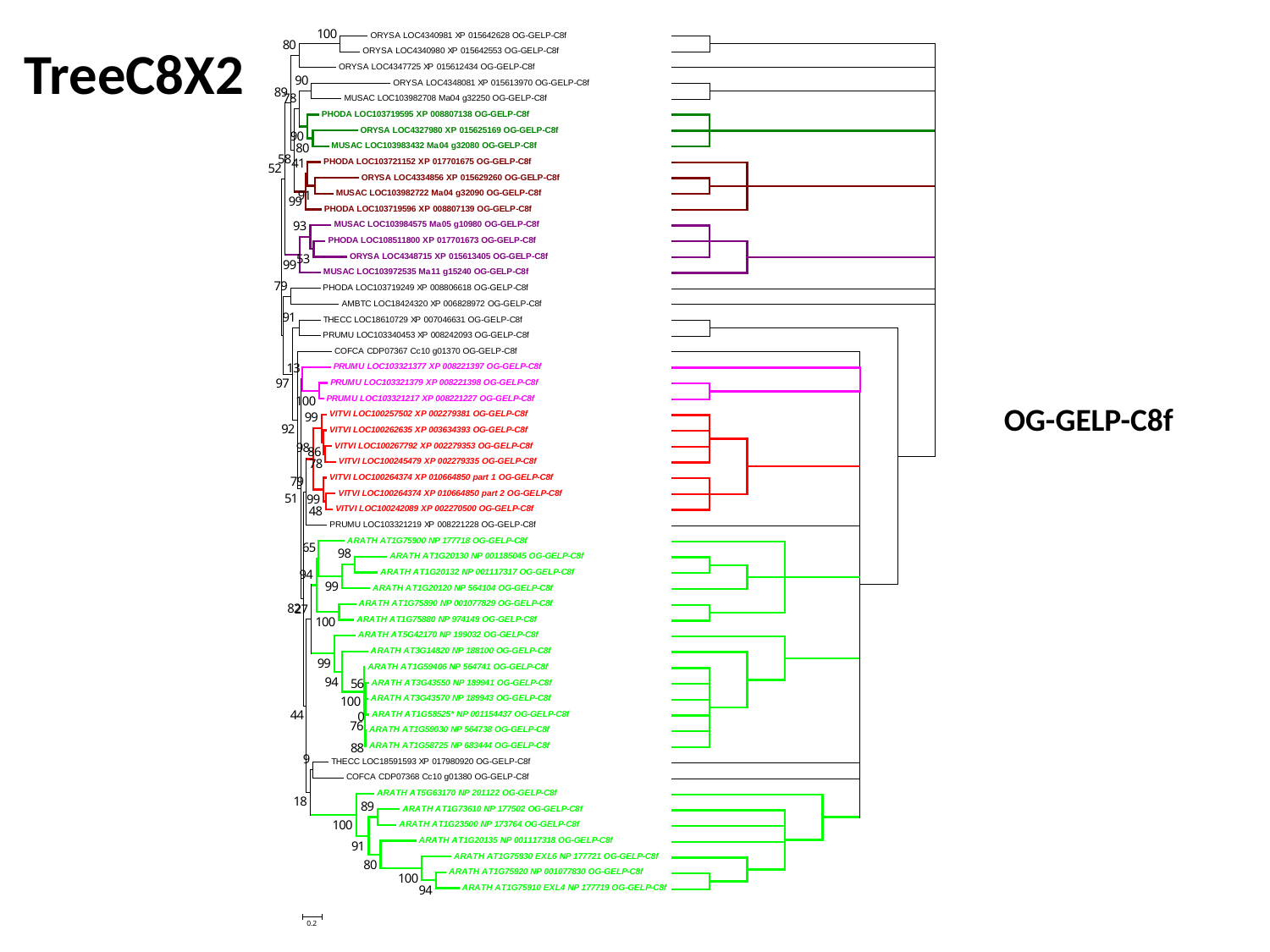

TreeC8X2
OG-GELP-C8f

### Slide 28
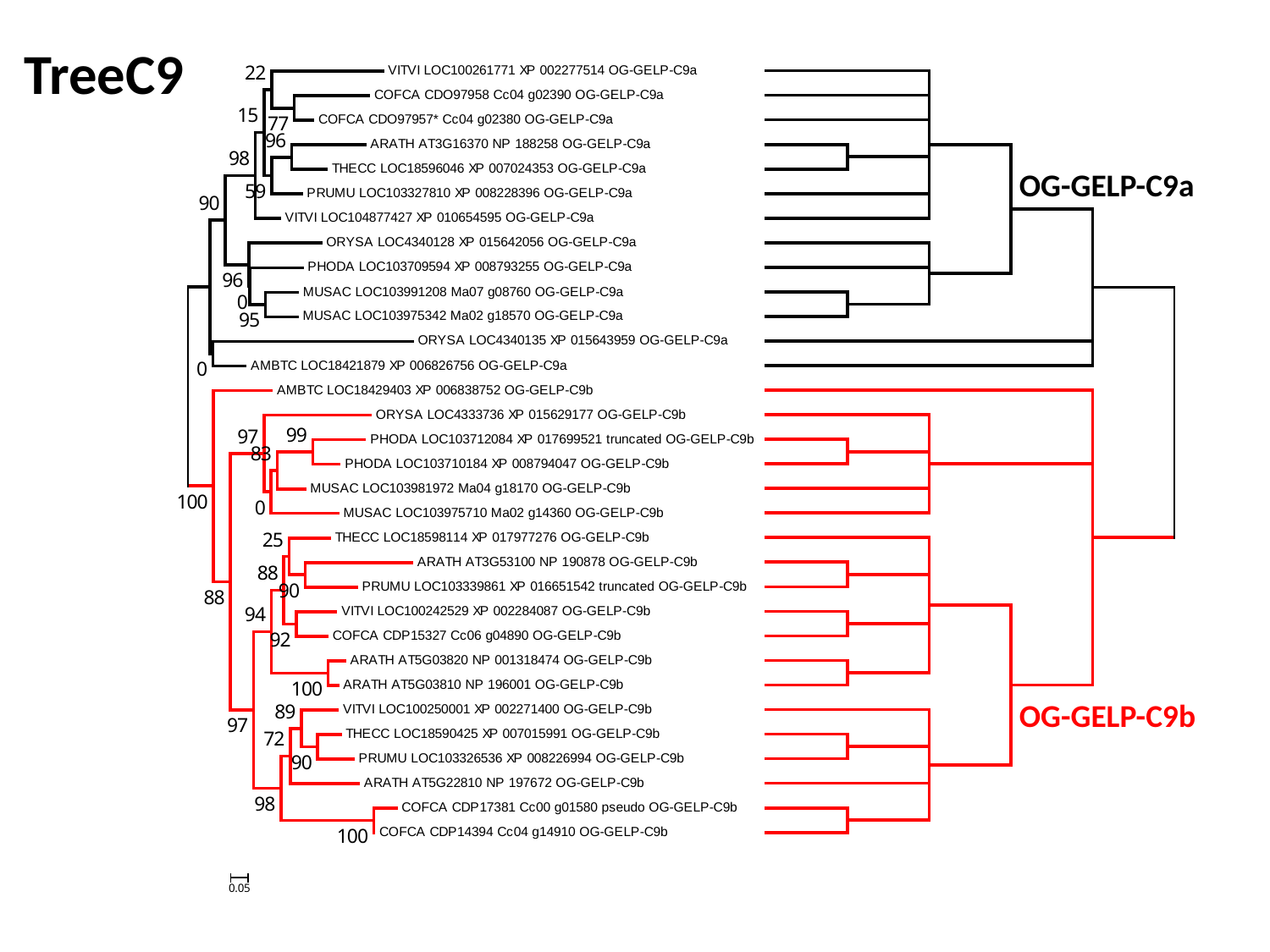

TreeC9
OG-GELP-C9a
OG-GELP-C9b

### Slide 29
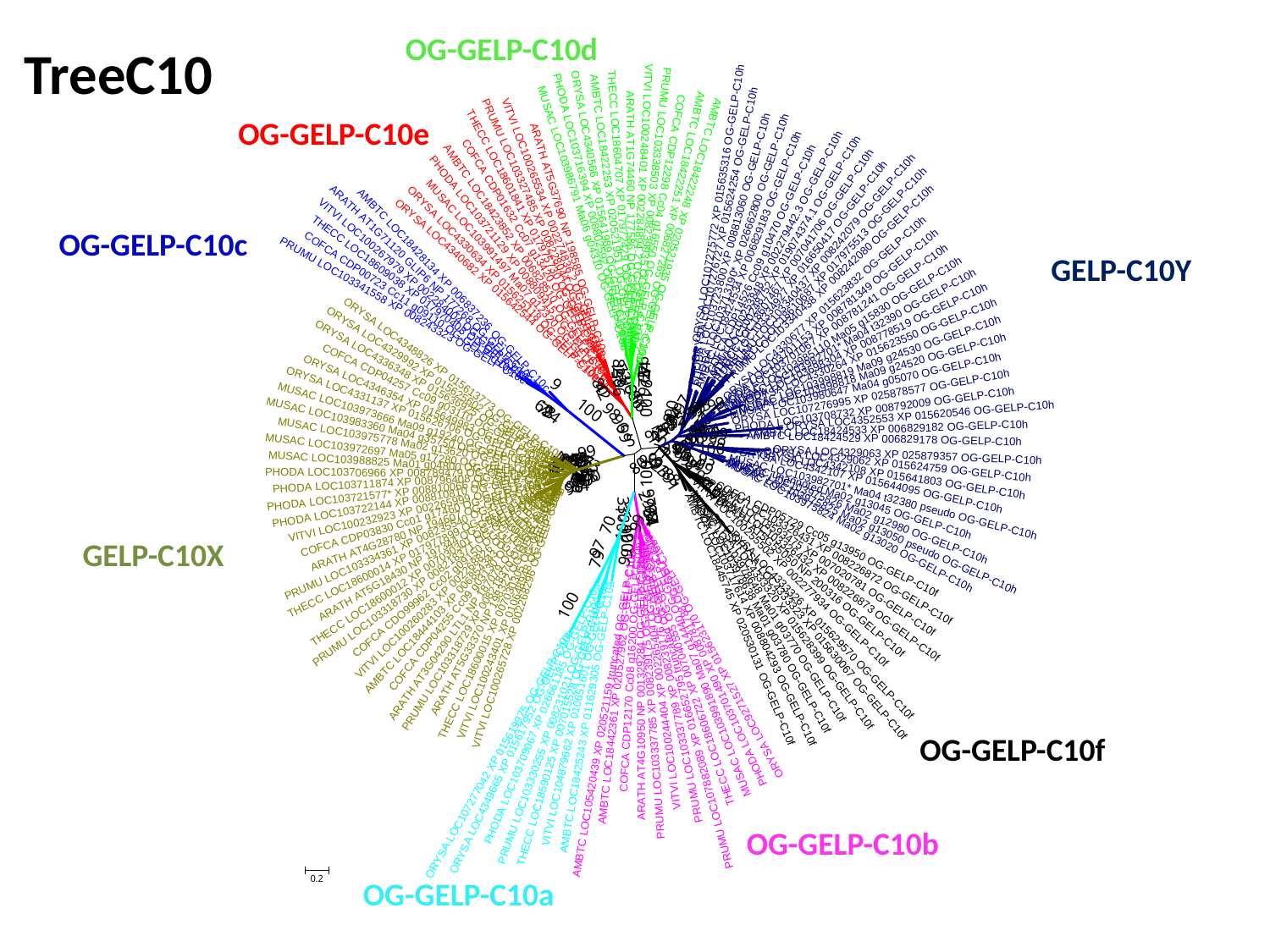

OG-GELP-C10d
TreeC10
OG-GELP-C10e
OG-GELP-C10c
GELP-C10Y
GELP-C10X
OG-GELP-C10f
OG-GELP-C10b
OG-GELP-C10a

### Slide 30
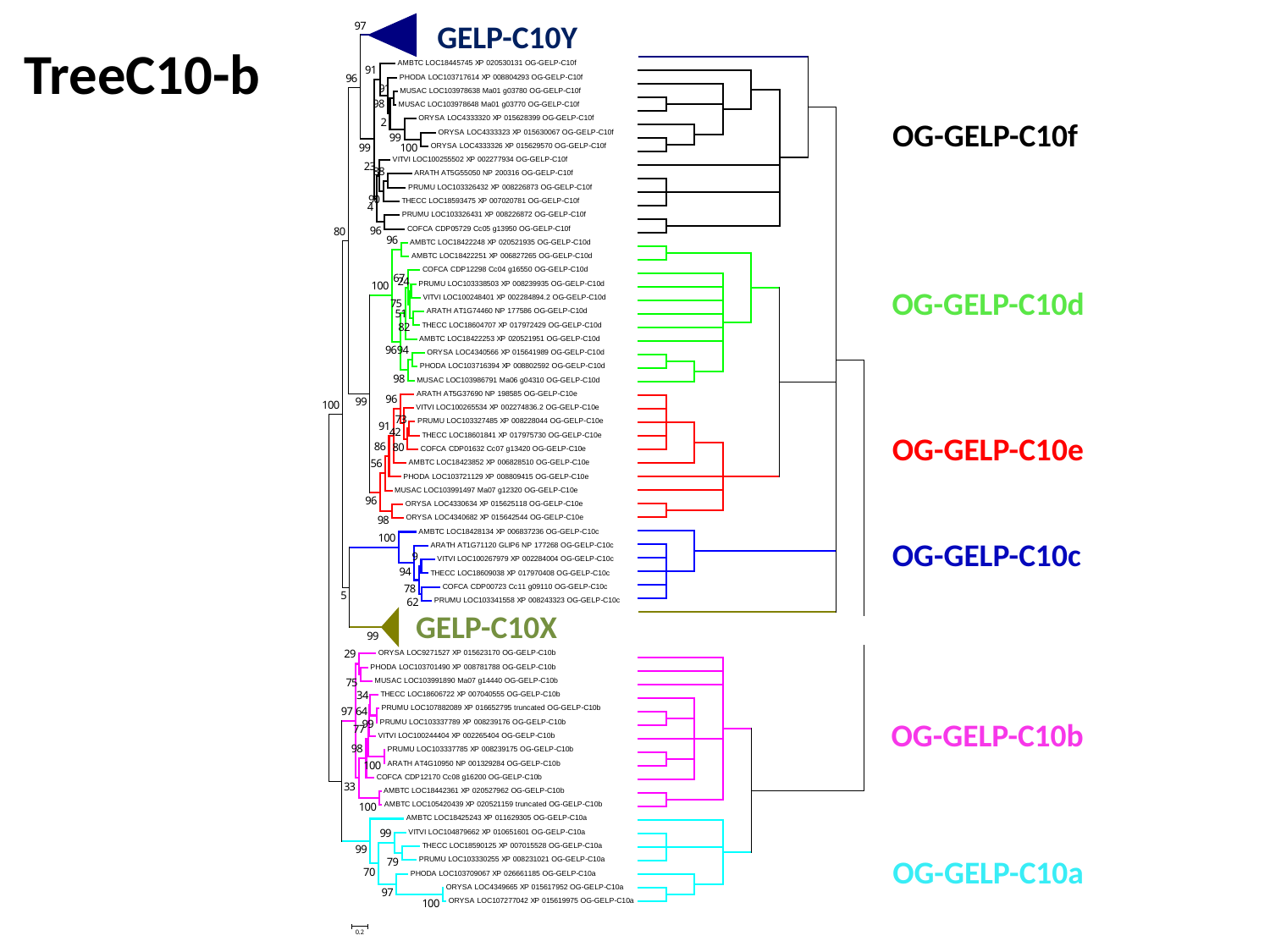

GELP-C10Y
TreeC10-b
OG-GELP-C10f
OG-GELP-C10d
OG-GELP-C10e
OG-GELP-C10c
GELP-C10X
OG-GELP-C10b
OG-GELP-C10a

### Slide 31
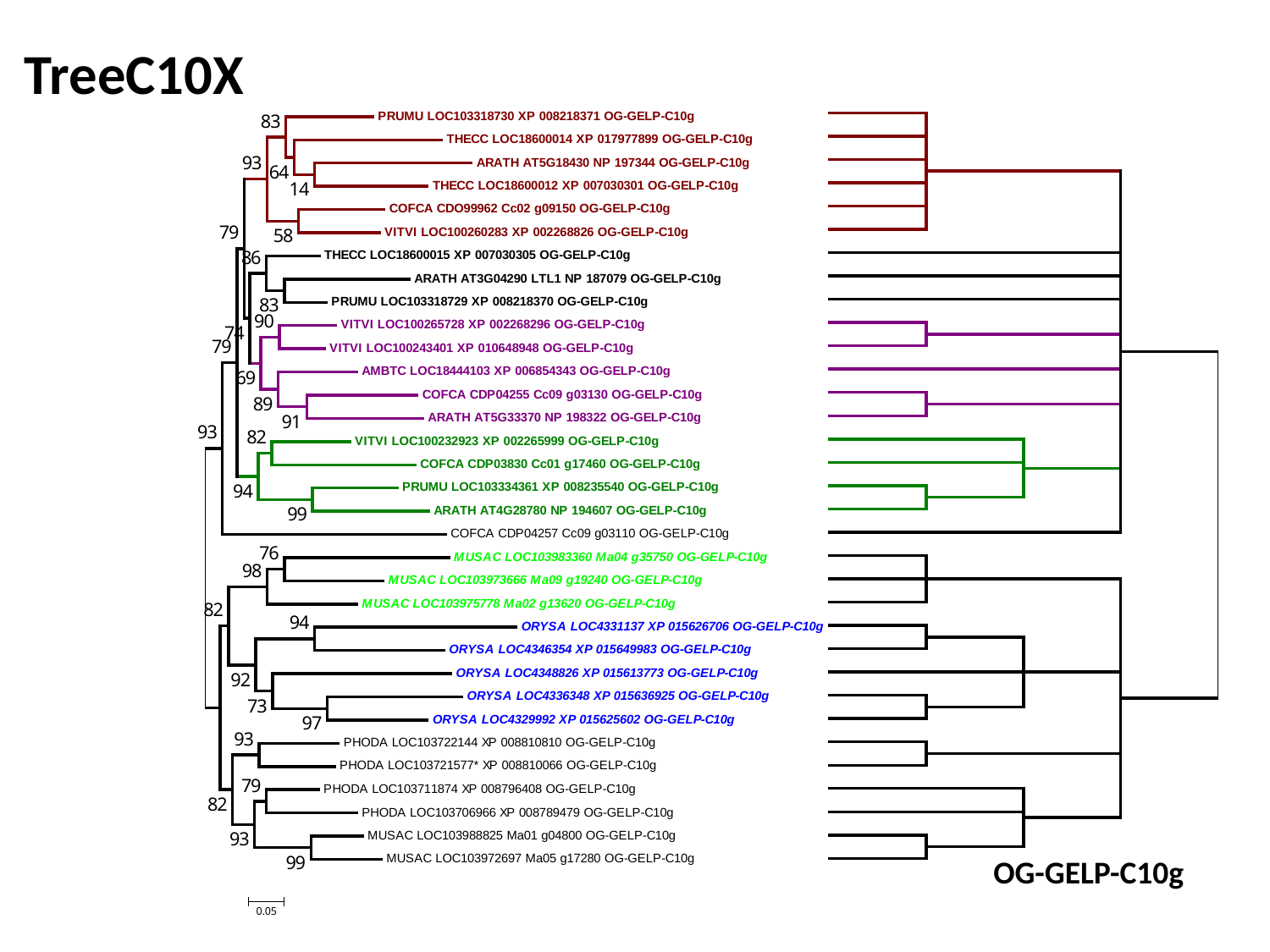

TreeC10X
OG-GELP-C10g

### Slide 32
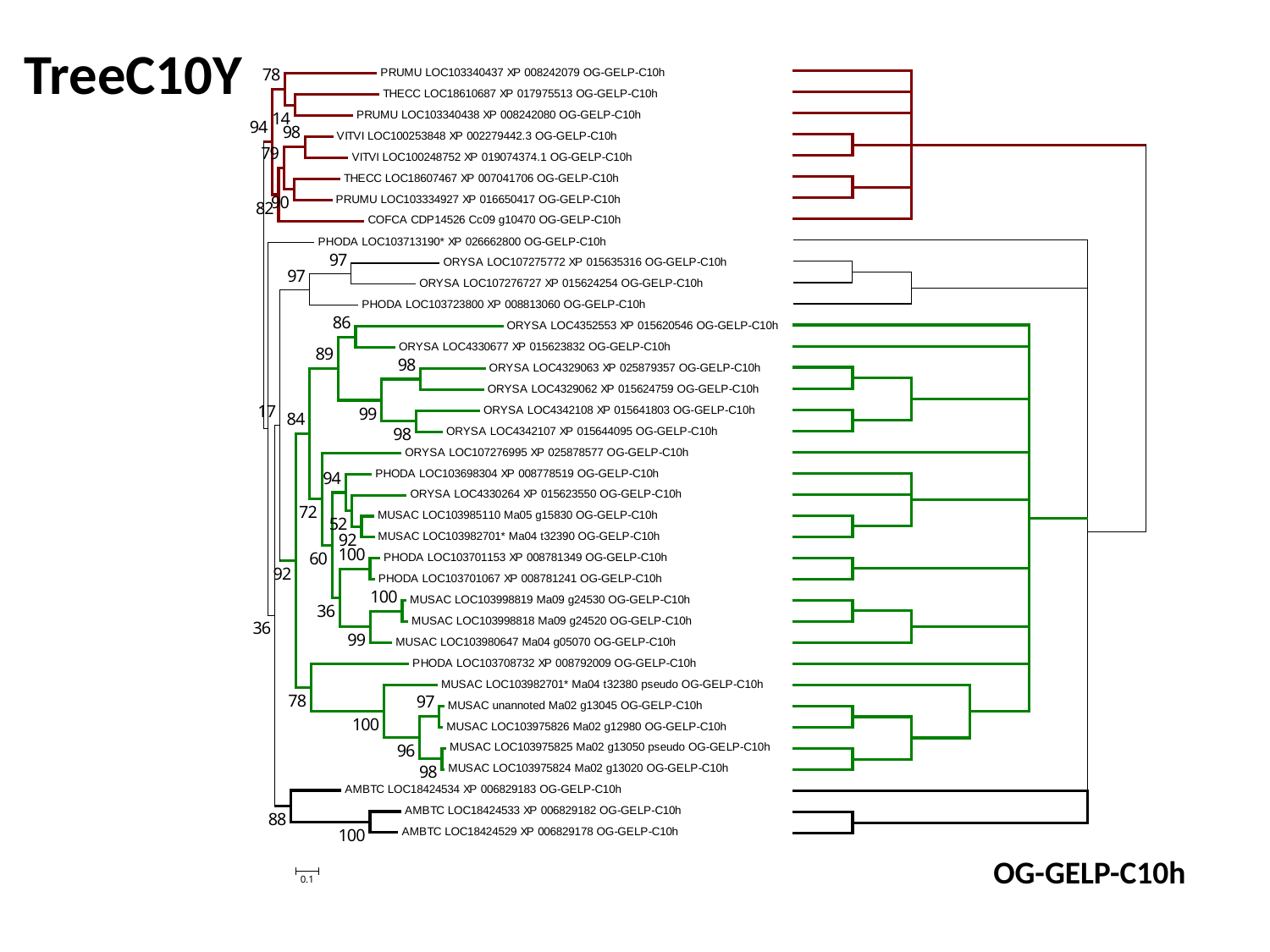

TreeC10Y
OG-GELP-C10h

### Slide 33
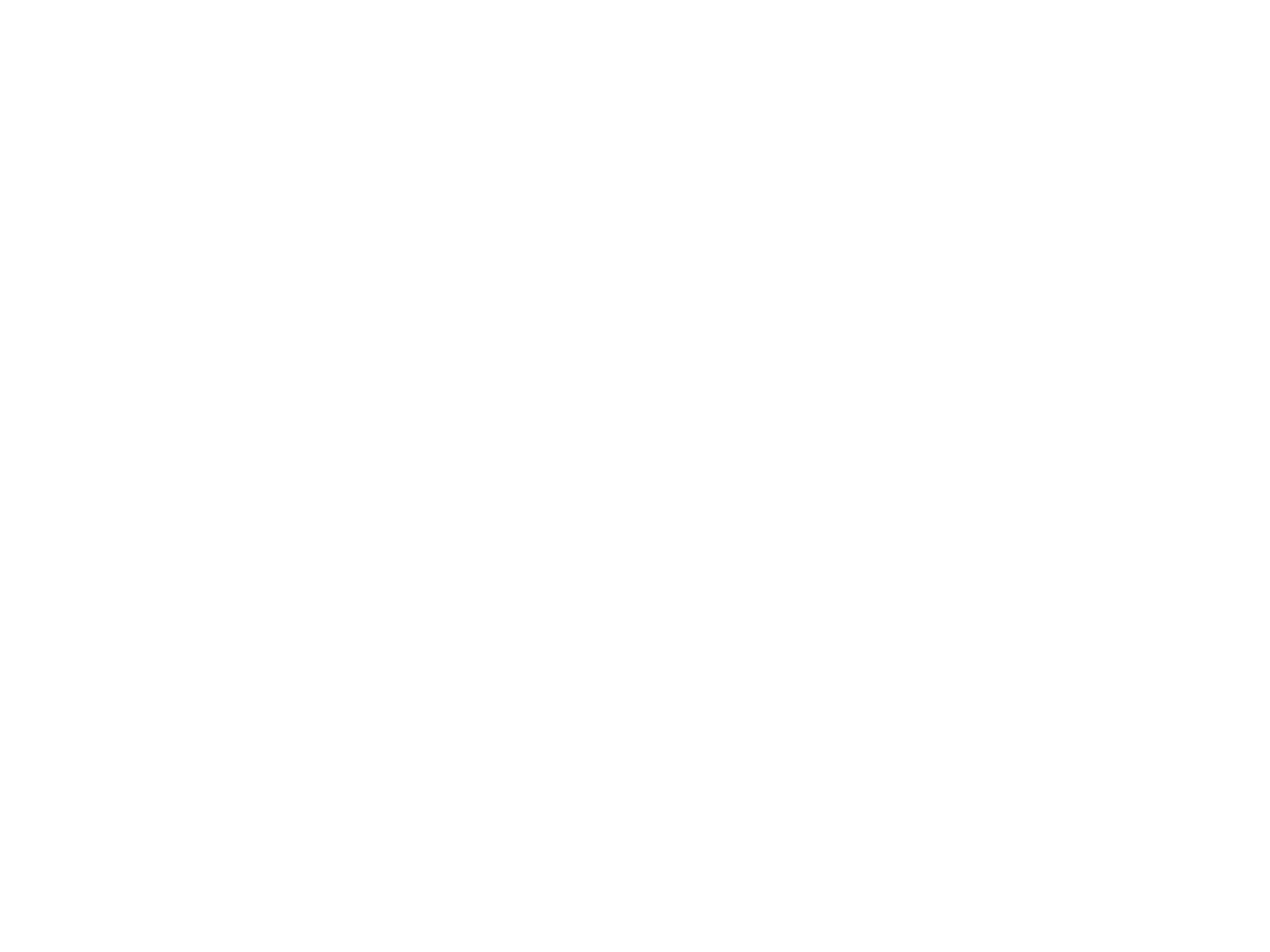
